# Effects of Long-Term Alcohol Consumption on Behavior in the P301S (Line PS19) Tauopathy Mouse Model

**DOI:** 10.1101/2022.07.12.499737

**Authors:** Christina M. Catavero, Annelise E. Marsh, Anthony M. Downs, Adonay T. Teklezghi, Todd J. Cohen, Zoe A. McElligott

## Abstract

Alcohol consumption and misuse remain prevalent public health issues with recent reports of increased heavy consumption in older populations in the United States. Several studies have identified alcohol consumption as a risk factor for developing multiple forms of dementia. The behavioral and psychological symptoms of dementia (BPSD), such as changes in affective behavior (e.g. anxiety), precede and coincide with cognitive decline. While many studies have characterized the intersection of alcohol use and affective behaviors, little is known regarding alcohol consumption and BPSD. This study characterizes the impact of long-term alcohol consumption on various behaviors in the P301S (line PS19) tauopathy mouse model. Male and female P301S and littermate control mice underwent two-bottle choice intermittent access to alcohol for sixteen weeks starting at 12 weeks of age. There were no significant differences in total ethanol consumption between wildtype and P301S mice of the same sex; however, drinking behavior differed among genotypes, and female mice of each genotype drank significantly more ethanol than males of each genotype. Following drinking studies, mice were run through a battery of behavioral tests during a period of forced abstinence, including approach/avoidance assays, social behavior tests, and memory and cognition tests. Across these tests we observed differences between groups due to genotype, alcohol history, and interactions between alcohol exposure and genotype. These differences were not always consistent between the sexes. In total, this study reveals significant alcohol-tauopathy interactions in subsequent behavior, which may have implications for understanding how alcohol may impact BPSD in conditions associated with tauopathy like Alzheimer’s disease and frontotemporal dementia.

## Introduction

Meta analyses suggest heavy alcohol consumption enhances risk for developing multiple forms of dementia, including the most common form, Alzheimer’s Disease (AD), as well as frontotemporal dementia (FTD) the most common form of dementia in persons under the age of 60 (Tyas, 2001; Ridley et al., 2013; Ilomaki et al., 2015; Gutwinski et al., 2018; Araujo et al., 2021). Over the past twenty years, the US population has seen a rise in the incidence of AD, as well as increased alcohol consumption in adults 50 years and older, particularly among women (Wilson et al., 2014; Breslow et al., 2017; Grucza et al., 2018; White, 2020; Araujo et al., 2021). Motivation to drink can vary among individuals and drinking phenotypes may be divided into “reward-drinkers” and “relief-drinkers” (Heinz et al., 2003; Glöckner-Rist et al., 2013; Mann et al., 2018). Rates of reward-drinking and relief-drinking among the elderly are unknown; furthermore, the relationship between different drinking phenotypes and aging, as well age-related disease like AD, is unclear, with conflicting reports of negative, positive, or no effect of alcohol consumption on AD progression (Tyas, 2001; Panza et al., 2012; Ilomaki et al., 2015; Heymann et al., 2016; Xu et al., 2017; Rehm et al., 2019; Kamal et al., 2020; Barnett et al., 2022). Therefore, it is difficult to assess correlation vs. causation, as well as directionality: does AD pathology drive increased alcohol consumption, or does increased alcohol consumption impact AD pathology?

Many individuals drink for the hedonic properties of alcohol (“reward-drinkers”). However, one hypothesis as to why individuals are driven to consume alcohol is to reduce negative affective states (“relief-drinkers”), and there are data suggesting individuals experiencing anxiety and depression are more likely to “self-medicate” with alcohol (Menary et al., 2011; Crum et al., 2013; Turner et al., 2018). Due to a variety of genetic and environmental factors, some individuals develop a physical dependence on alcohol, where upon cessation of consumption, profound withdrawal leads to seizures and the possibility of death. However, even in the absence of physical dependence, people experience a range of symptoms associated with alcohol withdrawal and may continue to consume alcohol despite negative consequences. In some individuals, alcohol consumption is driven by negative reinforcement to ameliorate the negative affective state associated with withdrawal (Koob and Le Moal, 1997, 2001; Wise and Koob, 2014; Koob, 2015). While dementia, particularly AD, is commonly associated with cognitive and memory deficits, mood changes are observed as well in which patients may feel depressed, anxious, fearful, irritable, and aggressive (Ferrazzoli et al., 2013; Todd, 2020). These symptoms are referred to as the behavioral and psychological symptoms of dementia (BPSD) and are estimated to occur in up to 90% of patients with AD, sometimes preceding cognitive decline (Bessey and Walaszek, 2019; Gottesman and Stern, 2019; Keszycki et al., 2019; Altomari et al., 2022). Regardless of motivation to drink, with the rise of alcohol consumption in older adults and the behavioral changes associated with AD, it is imperative to understand how alcohol use impacts AD pathology. The goal of this research is to lay groundwork for understanding the intersection of alcohol use and neurodegenerative diseases associated with tauopathy such as FTD and AD.

Transgenic mouse models expressing tauopathy and FTD- and AD-related pathology have been widely used to model cognitive impairments, however BPSD-like behaviors are not as well characterized. A review of over twenty mouse models of AD-related pathology (but notably not the P301S model) revealed recapitulation of many symptoms of AD such as aspects of aggression, sleep-wake disturbances, social withdrawal, and depressive-like and apathy-like behaviors; however, there is vast heterogeneity across changes in affective behavior, particularly approach-avoidance behaviors (Kosel et al., 2020; Pentkowski et al., 2021). This heterogeneity is likely driven by differences in the presentation and progression of pathology across models, as well differences in promoter-dependent expression of the transgene and the background strain (Kosel et al., 2020).

In the present study, we aimed to characterize the effect of long-term alcohol consumption on behavior in the humanized tauopathy mouse model P301S, the genetic mutation that has been associated with lethal early-onset FTD. We investigated this model due to the previously reported pattern of pathological tau spread throughout neural circuits implicated in emotion and reward, such as the locus coeruleus, hippocampus, and amygdala (Bugiani et al., 1999; Sperfeld et al., 1999; Morris et al., 2001; Lossos et al., 2003; Werber et al., 2003; Yoshiyama et al., 2007; Kang et al., 2020). We utilized the intermittent access (IA) to alcohol drinking paradigm, which is considered a model of non-dependent heavy alcohol consumption. IA results in persistent escalation of alcohol consumption due to repeated acute withdrawal periods within and between each weekly cycle of IA, and results in a greater escalation of alcohol consumption than continuous access (Hwa et al., 2011). In this study, P301S and wildtype littermate control male and female mice backcrossed on a congenic C57BL/6J background began drinking at three months of age for a duration of sixteen weeks to mimic chronic drinking across their adult life. 72 hours into a period of forced abstinence, the mice began a battery of behavioral assays lasting a period of one month. To our knowledge this is the first study of long-term alcohol consumption in the P301S mouse model, and furthermore, there have been few studies to examine how chronic alcohol consumption modulates behavior in aged mice. A previous study of long-term alcohol consumption in the 3x-Tg AD model found four months of alcohol consumption using the 2-bottle choice continuous access drinking paradigm impaired spatial memory, sensorimotor gating, and conditioned fear (Hoffman et al., 2019). Based on the results of Hoffman et al., we hypothesized long-term alcohol consumption in the P301S mice would exacerbate changes in behavior across a variety of approach/avoidance, memory, and cognitive tasks. We observed significantly divergent behavior across a number of assays driven by drinking history, genotype, and interactions between these two factors. Furthermore, we found differences in behavioral manifestations in male and female mice.

## Materials and Methods

### Subjects

For all experiments, adult male and female P301S (Jackson Labs) mice on a C57BL/6J background and wildtype littermate controls were bred in house and backcrossed over 20 generations. All animals in the study were 12-13 weeks old at the onset of drinking studies. Animals were singly housed and maintained on a reverse 12-hour light cycle with lights off 7:00am-7:00pm. Animals had *ad libitum* access to food and water, unless noted. All animal procedures were performed in accordance with the University of North Carolina at Chapel Hill’s institutional animal care and use committee’s regulations.

### Ethanol consumption paradigm

Intermittent Access (IA) to ethanol was performed in mouse home cages as previously described (Hwa et al., 2011). Briefly, experimental mice were singly-housed and provided access to an RHM 3000 diet (Prolab) one week prior to ethanol drinking. Mice then had access to a bottle of ethanol and water in their home cage on Monday, Wednesday, and Friday for 24-hour periods. On other days, they had access to two bottles of water. Bottles were rotated to prevent association of ethanol or water with a particular side of the cage. Control mice had access to two bottles of water every day. Mice underwent the intermittent access paradigm for sixteen weeks beginning at 12-13 weeks of age. Experimental mice had access to 3%, 6%, 10% (v/v) ethanol (unsweetened) weeks 1-3 of experiment (weekly at each dose), and access to 20% (v/v) ethanol (unsweetened) weeks 4-16 of experiment.

### Open Field (OF)

Mice were placed in an open field (50 cm x 50 cm) within a sound-attenuated chamber (Accuscan Instruments). Locomotor activity and position within the open field were measured for 30 min using infrared beam breaks. Light in the chamber was measured to be 40 lux. Data were analyzed using Accuscan software.

### Elevated Plus Maze (EPM)

Mice were placed in the center of the apparatus at the beginning of the test, and were given 5 min exploration time. Locomotor activity as well as position and duration within maze were monitored. The arms of the maze measure 77 cm x 77cm and the maze is elevated 74 cm above the ground. Light in the center of the EPM was measured to be 9.5 lux.

### Light-Dark box (LD box)

Mice were placed into the dark side of the apparatus (Med Associates) and latency to enter light side, time spent in light, and entries into light were monitored for 15 min using Ethovision version 15 (Noldus). Light in the chamber was measured to be 300 lux.

### Novelty-suppressed feeding

48 hours before testing, animals were allowed to consume a Froot Loop (Kellogg’s) in their home cage. All food was then removed from the home cage for 24 hours prior to the experiment. Mice were then placed in a corner of an open field at the center of which a single Froot Loop. Latency to feed was measured as the time required for the mouse to begin consuming the Froot Loop. If the mouse had not approached the Froot Loop after 10 min, it was removed from the open field and scored as 10 min. Immediately following the beginning of consumption, the mouse was returned to its home cage and allowed to freely consume Froot Loops for 10 min. If the mouse did not consume any Froot Loops in the home cage, it was excluded from this experiment. Light in the chamber was measured to be 10.5 lux.

### Free social interaction

Experimental mice and conspecific social target mice were habituated to the testing arena for 10 minutes 24 hours before testing. The testing arena consisted of a clear plastic mouse cage (70 square inches) with no bedding present. On test day, experimental mice and sex-matched social target mice (C57 albino, Jackson Labs) were placed into the testing arena and behavior was recorded for 10 min using an overhead camera. The duration of time that experimental mice spent actively engaging with the target mouse was measured by a blind observer. Active engagement is defined as the experimental mouse oriented towards the social target mouse (i.e., activity was not counted if the social target mouse was engaging in anogenital sniffing of the experimental mouse). The frequency and duration of socio-positive behaviors (nosing, anogenital sniffing, allogrooming, following, crawling) and agonistic behaviors (aggression, fighting) were recorded. Mice were excluded from this experiment when agonistic behaviors lasted longer than ten seconds/bout occurred to avoid injuries. Light in the chamber was measured to be 10.5 lux.

### Novel object recognition test (NOR)

The NOR test was performed based on a previously published protocol (Leger et al., 2013). Habituation: On day 1, mice were placed in the testing arena consisting of a clear plastic mouse cage (70 square inches) with no bedding present for ten minutes to habituate to environment. Training: On day 2, mice were placed in test chamber along with two plastic cubes of different colors for ten minutes. Testing: 24 hours later, on day 3, mice were placed in test chamber with one familiar plastic cube from the preceding day and one novel plastic shape for ten minutes. The time to reach 20 seconds of exploration among the two objects was recorded, as well as the preference ratio. Mice that did not reach 20s of exploration were excluded. Light in the chamber was measured to be 10.5 lux. Preference for novel object is reported on a scale 0-1.0: value > 0.5 = preference for novel object, value < 0.5 = preference for familiar object, value = 0.5 = equal preference.

### Conditioned fear

Training: On day 1, mice were placed in test chamber (Noldus Apparatus) contained in sound-attenuating boxes scented with 20% ethanol + 1% vanilla extract solution. The mice were allowed to explore the test chamber for 2 min baseline period and were then exposed to a 30-sec tone (80dB), followed by a 2s scrambled foot shock (0.6mA). Mice received two additional shock-tone pairings, with 80 sec between each shock-tone pairing.

Context-dependent recall: On day 2, mice were placed in original test chamber for a contextual learning test. Levels of freezing (immobility) in the absence of shock-tone pairings were determined across a 5-min session. Levels of freezing were measured throughout the entire session.

Cue-dependent recall: On day 3, mice were placed in a different sound-attenuated test chamber with walls and floor differing from the original test chamber, as well as a novel odor (0.5% acetic acid solution). Mice were placed in chamber and allowed to explore for 2 min baseline period, followed by acoustic stimulus (80dB tone) for a 3 min period. Levels of freezing before and during the stimulus were obtained.

### Statistical Analysis

Data are presented as mean ± SEM. * signifies main effect; @ genotype by treatment interaction. In figures, $ signifies genotype by time interaction; # treatment by time interaction. Significance is presented as *p<0.05, **p<0.01, ***p<0.001, ****p<0.0001. Two-way ANOVAs were used to compare genotype and treatment within each sex. Three-way ANOVAs were used to compare genotype, treatment, and time within each sex. Three-way ANOVAs were also used to compare sex, genotype, and treatment. Post-hoc Bonferroni corrections were used if a significant main effect was detected. Post-hoc Tukey corrections were used if a significant interaction was detected. Where ANOVAs were not applicable, the data was subjected to an unpaired t-test.

## Results

In order to examine the contribution of sex as a biological variable in addition the effects of ethanol treatment and the P301S mutant tau genotype, we utilized a 2 × 2 × 2 experimental design as depicted in **Fig. 1a, b**. This design yields four groups within each sex to study the interaction of ethanol-tau (MPE-<u>m</u>ale <u>P</u>301S <u>e</u>thanol, FPE-female <u>P</u>301S <u>e</u>thanol), ethanol alone (MWE- <u>m</u>ale <u>w</u>ildtype <u>e</u>thanol, FWE- female <u>w</u>ildtype <u>e</u>thanol), and tau alone (MPW- <u>m</u>ale <u>P</u>301S <u>w</u>ater, FPW- female <u>P</u>301S <u>w</u>ater). Wildtype mice consuming water were used as controls (MWW- <u>m</u>ale <u>w</u>ildtype <u>w</u>ater, FWW-female <u>w</u>ildtype <u>w</u>ater). **Fig. 1c** depicts the experimental timeline.

**Figure 1.**
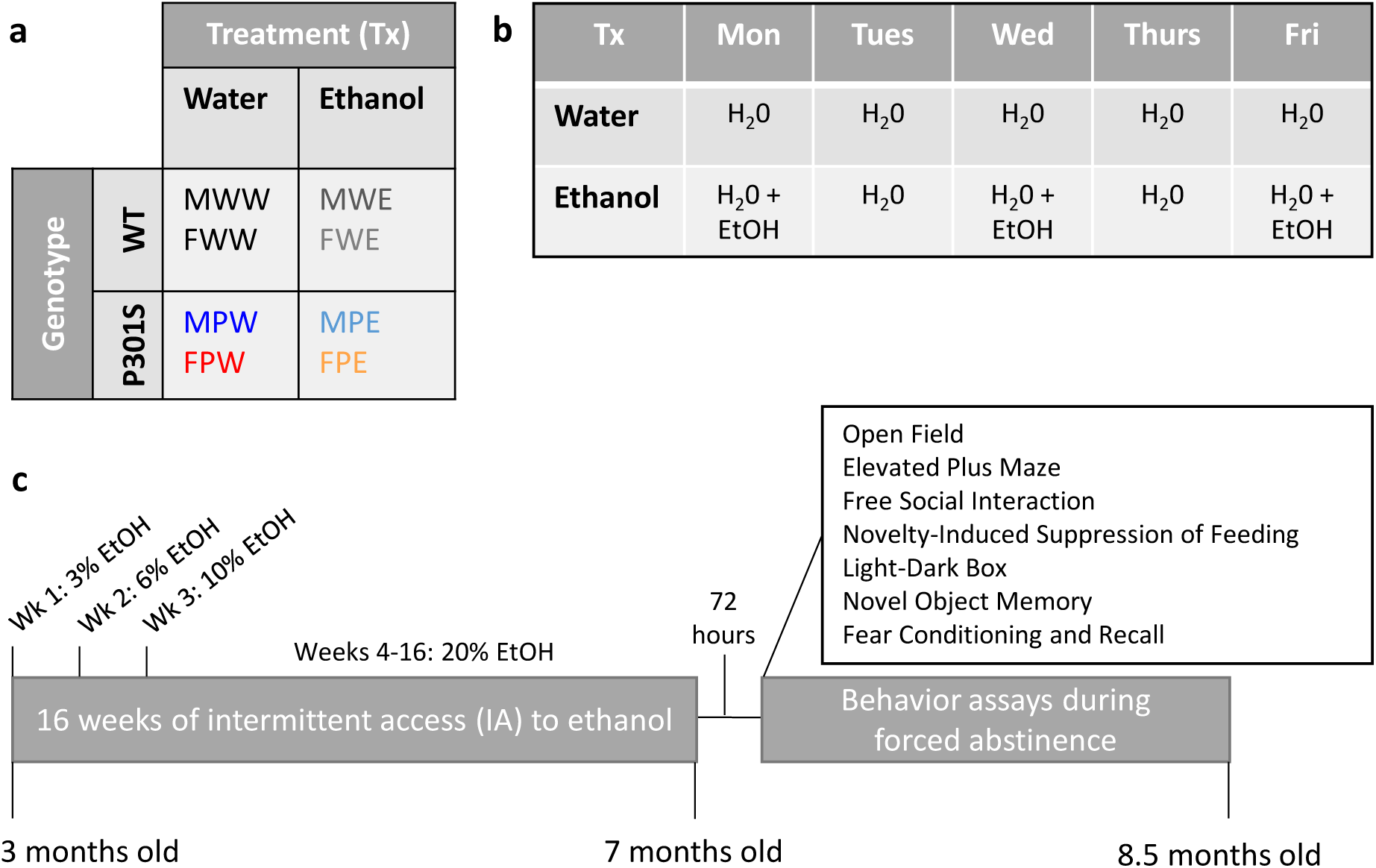
Experimental Design. **a) 2×2×2 Design.** Genotype (WT vs. P301S), treatment (Tx) (water vs. ethanol), sex (male vs. female). MWW = <u>M</u>ale <u>W</u>T <u>W</u>ater. MWE = <u>M</u>ale <u>W</u>T <u>E</u>thanol. MPW = <u>M</u>ale <u>P</u>301S Water. MPE = <u>M</u>ale <u>P</u>301S <u>E</u>thanol. FWW = <u>F</u>emale <u>W</u>T <u>W</u>ater. FWE = <u>F</u>emale <u>W</u>T <u>E</u>thanol. FPW = <u>F</u>emale <u>P</u>301S <u>W</u>ater. FPE = <u>F</u>emale <u>P</u>301S <u>E</u>thanol. **b) Treatment Paradigm**. Intermittent access to ethanol (IA). **c) Timeline**. Drinking behavior was observed among all groups for 16 weeks, ranging from 3 to 7 months of age. Behavior assays began three days after last drinking session and continued through 8.5 months of age. A minimum of 48 hours passed between each behavioral assay.

### Alcohol Consumption

We first examined the effect of drinking on changes in weight from baseline. Male P301S mice gained significantly less weight over the 16 weeks of drinking than their wildtype littermates (**Fig. 2a: Male change in weight from baseline**: Three-way ANOVA. Genotype x Week, F (15,1215) = 4.324, p <0.0001****; Genotype, F (1, 81) = 3.720, p = 0.0743; Week, F (1, 81) = 0.0000, p = 0.9928; post-hoc Tukey’s did not further reveal significant differences). There was a significant treatment by week interaction in female mice, such that female mice that consumed ethanol gained more weight compared to genotype-matched controls (**Fig. 2b: Female change in weight from baseline:** Three-way ANOVA. Treatment x Week, F (15,1245) = 2.225, p <0.0045**; Treatment, F (1, 83) = 0.8602, p = 0.3564; Genotype, F (1, 83) = 0.6849, p = 0.4103; post-hoc Tukey’s did not further reveal significant differences).

**Figure 2.**
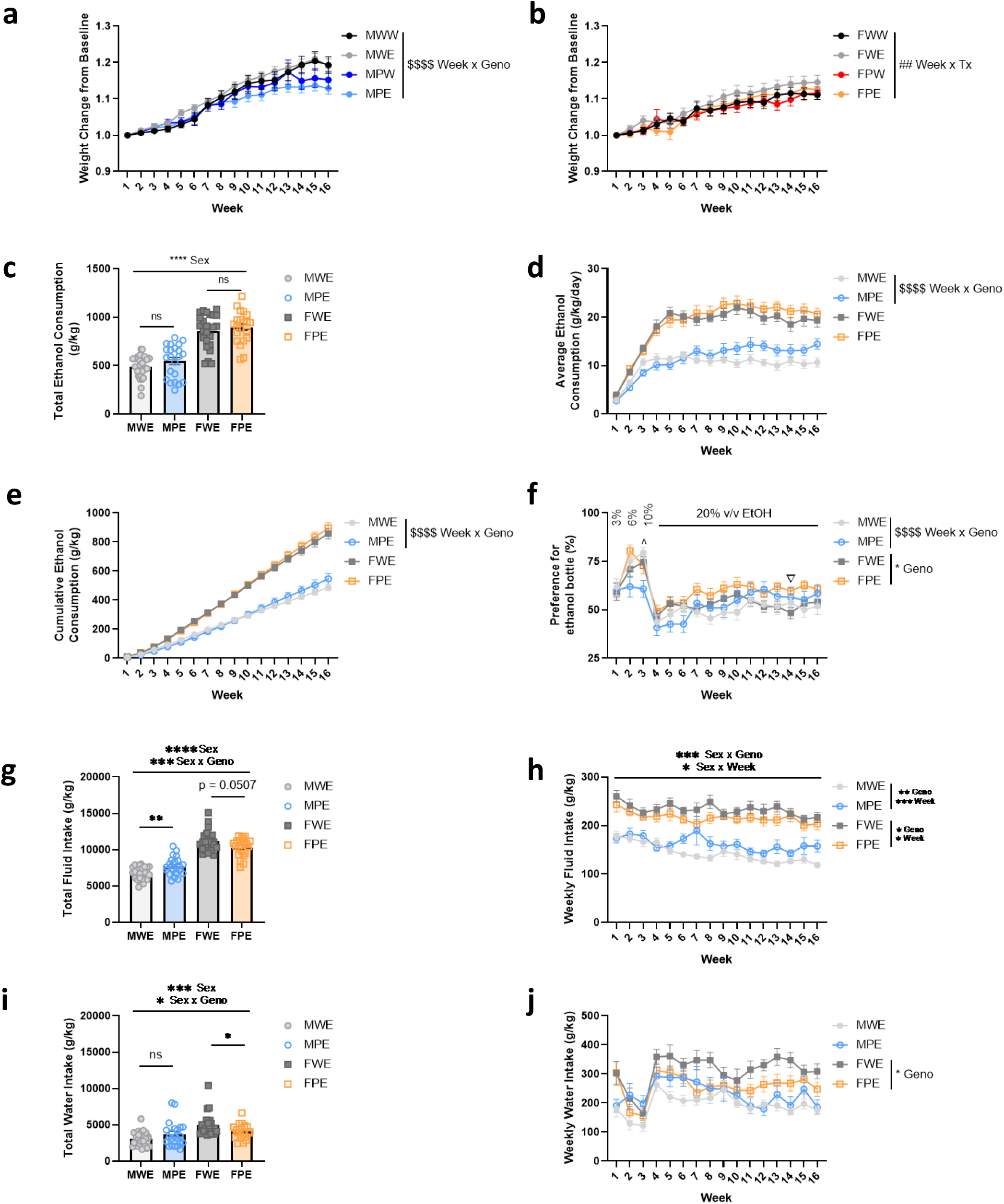
Drinking behavior. **(a)** Male weight change from baseline **(b)** Female weight change from baseline **(c)** Total ethanol consumption (g/kg) (**d)** Average daily ethanol consumption (g/kg/day) **(e)** Cumulative ethanol consumption (g/kg) **(f)** Preference for ethanol bottle (%) **(g)** Total fluid intake during drinking paradigm (g/kg) **(h)** Weekly fluid intake during drinking paradigm (g/kg) **(i)** Total water intake during drinking paradigm (g/kg) **(j)** Weekly water intake during drinking paradigm (g/kg). MWW = Male WT Water. MWE = Male WT Ethanol. MPW = Male P301S Water. MPE = Male P301S Ethanol. FWW = Female WT Water. FWE = Female WT Ethanol. FPW = Female P301S Water. FPE = Female P301S Ethanol. MWW n = 20, MWE n = 26, MPW n = 17, MPE n = 22, FWW n = 21, FWE n = 24, FPW n = 21, FPE n = 21.

Ethanol drinking behavior in the P301S mouse line has not been previously described. Over the course of sixteen weeks of IA, there are no significant differences in total ethanol consumption between wildtype and P301S mice within each sex; however, female mice drink more than experimental condition-matched males. **(Fig. 2c: Total ethanol consumption (grams ethanol/kilograms body weight):** Two-way ANOVA, Sex, F (1,89) = 118.5, p < 0.0001****. Post-hoc Bonferroni multiple comparison correction, ns. Post-hoc planned unpaired t-tests were performed due to differences in ethanol consumption: males, t = 1.435, df = 46, p = 0.1580, ns; females, t= 0.7542, df = 43, p = 0.4548, ns.) Additionally, when examined by week, male, but not female, mice exhibited a significant genotype by time interaction in both average daily consumption plotted by week as well as cumulative weekly consumption; however, comparisons between the groups did not survive post-hoc tests (**Fig. 2d: Average ethanol consumed (grams ethanol/kilograms bodyweight/day):** Two-way ANOVA males, genotype x time, F (15, 690) = 4.030, p < 0.0001****, genotype, F (1, 46) = 1.996, p = 1.645; post-hoc Tukey test did not further reveal significant differences; Two-way ANOVA females, genotype x time F (15, 645) = 0.7295, p = 0.7554, ns; genotype, F (1,43) = 0.5658, p = 0.4560, ns. **Fig. 2e: Cumulative ethanol consumed (grams ethanol/kilograms bodyweight/week):** Two-way ANOVA males, genotype x week, F (15, 690) = 2.969, p = 0.0001***; genotype, F (1, 46) = 0.2137, p = 0.6461, ns; post-hoc Tukey test did not further reveal significant differences; Two-way ANOVA females, genotype x week, F (15, 645) = 0.6929, p = 0.7929, ns; genotype, F (1,43) = 0.1408, p = 0.7093). Examination of preference for ethanol bottle revealed a significant genotype by time interaction in male mice and a main effect of genotype in female mice (**Fig 2f: Preference for ethanol bottle (%):** Two-way ANOVA males, genotype x week, F (15,690) = 3.183, p < 0.0001****; genotype, F (1, 46) = 0.0001, p = 0.9907, ns; post-hoc Tukey test: Week 3, p = 0.0170*; Two-way ANOVA females, genotype x week, F (15, 6450 = 1.230, p = 0.2437, ns; genotype, F (1,43) = 4.176, p < 0.0472*; post-hoc Tukey test: Week 14: p = 0.0359*).

We next examined total fluid intake to assess differences across the ethanol-drinking groups in fluid homeostasis and/or satiation. Total fluid intake is a measure of the amount of fluid consumed from the water bottle and the ethanol bottle; this measure takes into account the water in the ethanol bottle, in contrast to the measures of ethanol consumed in **Fig. 2c-e** which only account for the ethanol within the solution. Due to sex differences in ethanol drinking among males and females, we analyzed total fluid intake and water intake by two-way ANOVA to assess for sex differences, and planned unpaired t-tests within sex for a more sensitive analysis. Female mice consume significantly more fluid than males, and there is a sex by genotype interaction such that there is a significant increase in fluid intake among P301S males compared to WT, and a trend towards a decrease among female P301S compared to WT (**Fig. 2g: Total fluid intake (grams fluid/kilograms body weight):** Two-way ANOVA, both sexes: sex x genotype, F (1, 89) = 12.65, p = 0.0006**; sex, F (1, 89) = 192.3, p<0.0001****; Bonferroni’s multiple comparison’s test: males p = 0.0123*, females p = 0.0563. Unpaired t-test, males t = 3.174, df = 46, p = 0.0027**; Unpaired t-test, females t = 2.010, df = 43, p = 0.0507). When examining fluid intake by week, there are significant main effects of genotype for both sexes **(Fig. 2h: Weekly fluid consumption (grams fluid/kilograms body weight):** Two-way ANOVA males, genotype F (1,46) = 9.567, p = 0.0034**; Two-way ANOVA females, genotype F (1,43) = 4.297, p = 0.0442*. Post-hoc testing did not reveal further significant differences).

We then examined total water intake to assess there are differences in water homeostasis. There is a significant sex by genotype interaction and main effect of sex; however, due to differences in ethanol and total fluid consumption between males and females, we further analyzed water intake using planned unpaired t-tests within each sex which only revealed a difference in females but not males (**Fig. 2i: Total water intake (grams water/kilograms body weight):** Two-way ANOVA, both sexes: sex x genotype, F (1, 89) = 6.916, p = 0.0101*; sex, F (1, 89) = 16.53, p<0.0001****. Unpaired t-test, males t = 1.507, df = 46, p = 0.1386; Unpaired t-test, females t = 2.220, df = 43, p = 0.0317*). Additionally, when examined by week, there is a main effect of genotype on water intake among females but not males **(Fig. 2j: Weekly water intake (grams water/kilograms body weight):** Two-way ANOVA males, F (1, 46) = 2.271, p = 0.1386 ; Two-way ANOVA females, F (1, 43) = 4.938, p = 0.0316*; post-hoc testing did not reveal further significant differences).

Together these data suggest subtle differences in drinking behavior across genotypes within each sex. While there are no differences in total ethanol consumption among female wildtype and P301S mice, the P301S females consume significantly less water, driving a trend towards decreased fluid intake, and significant increase in preference for ethanol consumption. Among male mice, while there was no difference in total ethanol consumption between wildtype and P301S, examining ethanol consumption and preference by week revealed significant genotype by week interactions, such that MPE mice had greater total and weekly fluid intake compared to MWE but no difference in water intake. Together these data suggest subtle changes in ethanol drinking behavior among P301S mice over sixteen weeks of IA. However, due to profound sex differences in total ethanol consumption, water intake, and fluid intake – and thus total alcohol dosage over the 16 weeks of drinking – males and females were analyzed separately in subsequent behavioral tests.

### Approach/Avoidance Assays

Given tau pathology impacts brain regions implicated in appetitive and aversive behaviors and alcohol misuse is comorbid with anxiety disorders, we conducted a series of behavioral tests to broadly measure approach/avoidance behavior in mice as a proxy of aspects of BPSD-like behavior (Prut and Belzung, 2003; Gallagher et al., 2018; Turner et al., 2018; Kosel et al., 2020; Pentkowski et al., 2021). By using multiple tests, we are able to capture the nuance of these behaviors in rodents in novel and brightly lit (aversive to mice) or more dim environments, which beyond approach/avoidance also includes inhibitory/disinhibitory behavior. Additionally, by measuring distance traveled and/or velocity in several of these tests, we are able to observe locomotor behavior, and how this behavior is affected by environmental conditions of the maze and the lighting.

#### Open Field

In the open field (OF) test, P301S males traveled further compared to wildtype in terms of (**Fig. 3a. Males Total Distance Traveled**, two-way ANOVA: genotype x treatment: F (1, 38) = 6.032, p < 0.0187****; treatment F (1, 38) = 0.02571, p = 0.8735, genotype F (1, 38) = 21.24, p <0.0001****; post-hoc Tukey’s test: MWW vs. MPE, p = 0.0163*, MWE vs. MPW, p = 0.0090**, MWE vs. MPE, p < 0.0001****), as well as when examining locomotion in 10 minute bins across the 30 minute test as a measure of both locomotion and habituation (**Fig. 3b. Males Distance Traveled in Ten Minute Time Bins**, three-way ANOVA: genotype x treatment F (1, 37) = 4.169, p = 0.0484*; time x treatment F (2, 74) = 5.290, p = 0.0071**, genotype F (1, 37) = 22.56, p < 0.0001****; post-hoc Tukey’s test: 0-10: MWE vs. MPW, p = 0.0046**; 0-10: MWE vs. MPE, p = 0.0057**; 10-20: MWE vs. MPE, p = 0.0034**; 20-30: MWW vs. MPE, p = 0.0223*; MWE vs. MPW, p = 0.0470*; MWE vs. MPE, p = 0.0002***. **Fig. 3c. Males Average Velocity in Ten Minute Bins**, three-way ANOVA: genotype F (1, 37) = 18.13, p = 0.0001***; treatment F (1,37) = 0.3258, p = 0.5716; treatment x time F (2, 74) = 3.011, p = 0.0320*’ post-hoc Tukey’s test: 10-20: MWE vs. MPW, p = 0.0458*; 10-20: MWE vs. MPW p = 0.0494*, MWE vs. MPE p = 0.0096**; 20-30: MWE vs. MPE p = 0.0017**). While there was no difference in total distance traveled among female groups (**Fig. 3d. Females Total Distance Traveled**, two-way ANOVA: genotype F (1,35) = 0.5502, p = 0.4632; treatment F (1, 35) = 0.0139, p = 0.9069), P301S females sustained increased locomotion throughout the 30 minute test suggesting they did not habituate to the arena compared to WT (**Fig. 3e. Females Total Distance Traveled in Ten Minute Time Bins**, three-way ANOVA: time x genotype: F (2, 70) = 0.01387, p = 0.0061**; genotype F (1, 35) = 0.5502, p = 0.4632; treatment F (1, 35) = 0.0139, p = 0.9069; post-hoc testing did not reveal further significant differences). Additionally, P301S female mice exhibited greater average velocity and there was a significant alcohol-genotype interaction (**Fig. 3f. Females Average Velocity in Ten Minute Time Bins**, three-way ANOVA: time x genotype: F (2, 70) = 4.147, p = 0.0199*; genotype x treatment: F (1, 35) = 7.513, p = 0.0038**; post-hoc Tukey’s test: 10-20: FWW vs. FPW p = 0.0098**, FWE vs. FPW, p = 0.0257*; 20-30: FWW vs. FPW, p = 0.0198*; FEW vs. FPW p = 0.0452*). The time spent in zones of the OF arena was also analyzed to examine changes in avoidance behavior (**Fig. 3g-l**). There was a trend towards male P301S mice spending less time in the corners of the maze compared to WT (**Fig 3h. Males Time Spent in Corner**, two-way ANOVA: genotype F (1,38) = 3.905, p = 0.0554). However, there were no significant differences in time spent in the center or surround for any of the male groups (**Fig. 3g, i**), nor any significant differences in time spent in any of the zones among the female groups (**Fig 3j-l)**.

**Figure 3.**
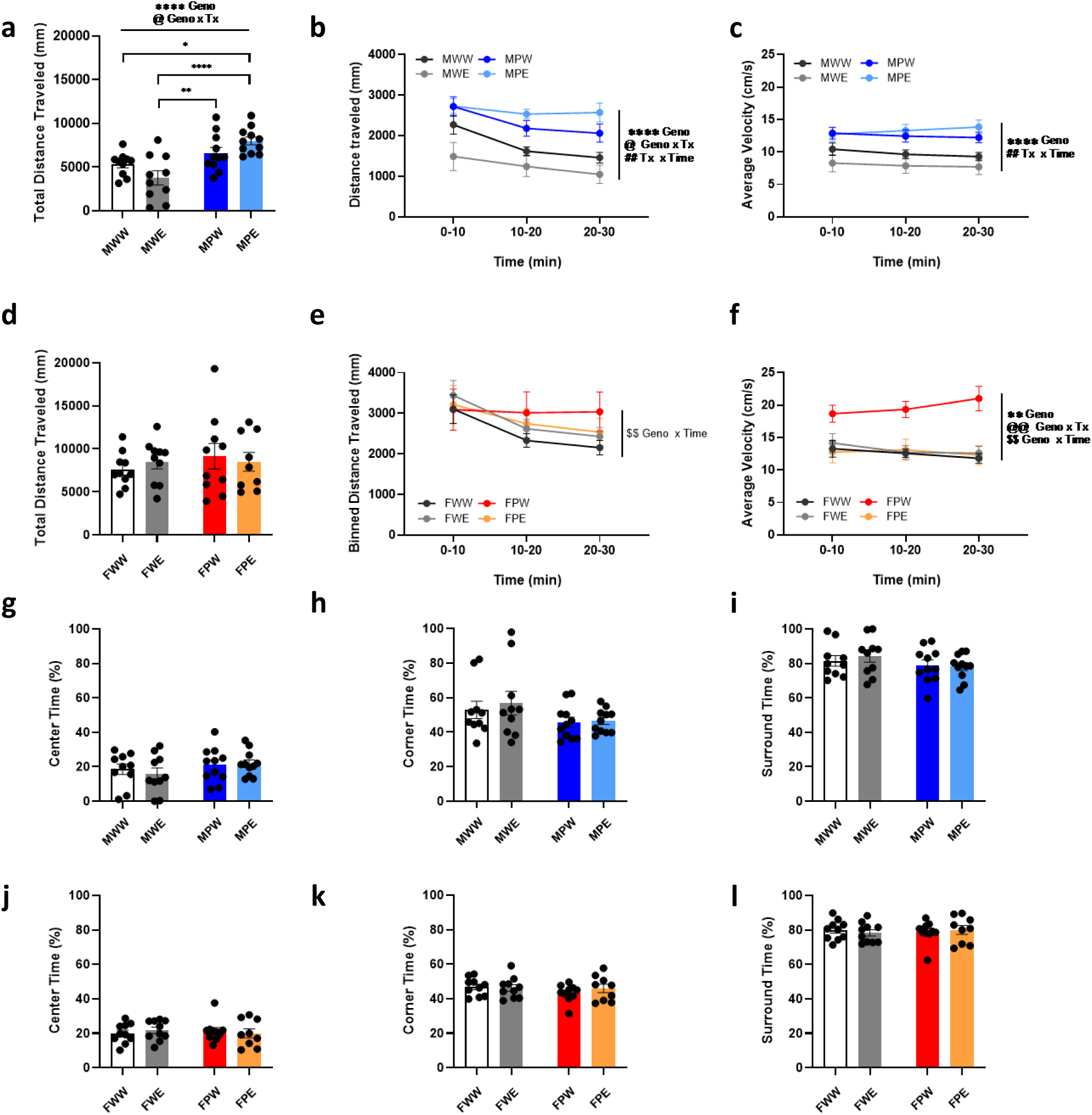
Open Field Test. **(a)** Males total distance traveled (mm). **(b)** Males total distance traveled in ten minute time bins (mm). **(c)** Males average velocity in ten minute time bins (cm/s). **(d)** Females total distance traveled (mm). **(e)** Females total distance traveled in ten minute time bins (mm). **(f)** Females average velocity in ten minute time bins (cm/s). **(g, j)** Time in center of open field (%). **(h, k)** Time in corners of open field (%). **(i, l)** Time in surround of open field (%).

### Elevated Plus Maze

In the elevated plus maze (EPM), conducted in a low light environment, male P301S mice spent significantly more time in the open arms compared to WT (**Fig. 4a, Males Time in open arms**, two-way ANOVA: genotype F (1, 38) = 8.005, p = 0.0074**; treatment F (1,38) = 0.6890, p = 0.4117) as well as less time in closed arms (**Fig. 4c, Males Time in closed arms**, two-way ANOVA: genotype F (1, 38) = 4.164, p = 0.0483*; treatment F (1, 38) = 0.4307, p = 0.5156; Post hoc multiple comparisons revealed P301S males exposed to ethanol spent less time in closed arms (**Fig. 4c**, Bonferroni, WT-P301S, water p = 0.8884, ethanol p = 0.0379*), and more time in the open arms compared to WT males exposed to ethanol (**Fig. 4a**, Bonferroni, WT-P301S, water p = 0.1952, ethanol p = 0.0504).

**Figure 4.**
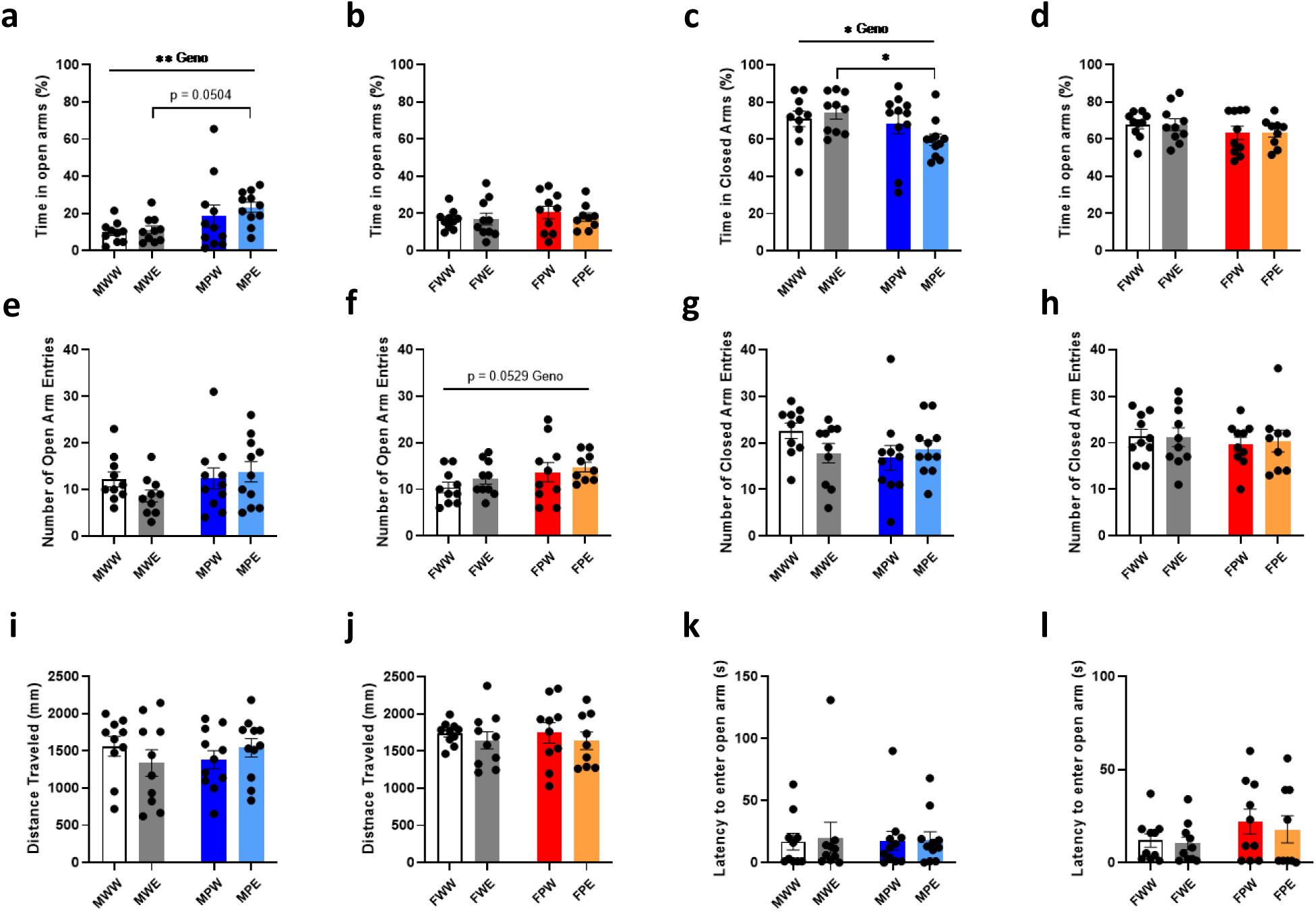
Elevated Plus Maze. **(a, b)** Time in open arms (%). **(c, d)** Time in closed arms (%). **(e, f)** Number of open arm entries. **(g, h**) Number of closed arm entries. **(I, j)** Distance traveled (mm) **(k, l)** Latency to enter open arm (s).

There were no significant differences among female groups regarding time spent in the open and closed arms (**Fig. 4b, d**), however there was a strong trend towards P301S females entering the open arm more frequently compared to WT (**Fig. 4f, Females Number of open arm entries**, two-way ANOVA: genotype F (1, 35) = 4.013, p = 0.0529). We found no significant difference in distance traveled, closed arm entries, or latency to enter an open arm in the maze for male and female mice (**Fig. 4d, h, i-l**, Two-way ANOVA).

### Novelty-Induced Suppression of Feeding Test

The NSF test is used to assess approach/avoidance behavior in a novel, brightly lit environment. Latency to eat is a proxy for anxiety-like behavior measured as motivation to approach food in a novel environment. In male mice there were no significant differences in latency to feed (**Fig. 5a, Males latency to feed**, two-way ANOVA: genotype x treatment: F (1, 29) = 1.342, p = 0.2561 ns; treatment F (1, 29) = 0.04025, p = 0.8424 ns; genotype F (1, 29) = 0.6055, p = 0.4428 ns). Female P301S mice took significantly longer to feed (**Fig. 5b, Females latency to feed**, two-way ANOVA: genotype x treatment: F (1, 40) = 0.5293, p = 0.4712 ns; treatment F (1, 40) = 1.709, p = 0.1986 ns; genotype F (1, 40) = 4.272, p = 0.0453*). Post-hoc Bonferroni’s multiple comparisons test indicated no significant differences, although there was a trend for a difference in the water consuming animals (WT Females – P301S Females: water: F (1,40), p = 0.0997; ethanol F (1,40), p = 0.7199). Post-test consumption is assessed as a control measure to ensure there are no differences in baseline hunger between groups. There are no significant differences in post-test home cage consumption in male and female mice. (**Fig. 5c, Males, 5d Females Post-test homecage consumption**, Males: two-way ANOVA: genotype x treatment: F (1, 29) = 0.6905, p = 0.4128 ns; treatment F (1, 29) = 1.262, p = 0.2705 ns; genotype F (1, 29) = 2.156, p = 0.1528 ns; Females: two-way ANOVA: genotype x treatment: F (1, 40) = 1.016, p = 0.3194 ns; treatment F (1, 40) = 0.7811, p = 0.3821 ns; genotype F (1, 40) = 0.4281, p = 0.5167 ns). Mice that did not feed in their home cage on days 1 or 3 were excluded from the data set. Together thees data suggest female P301S mice express more avoidance behavior in the NSF test.

**Figure 5.**
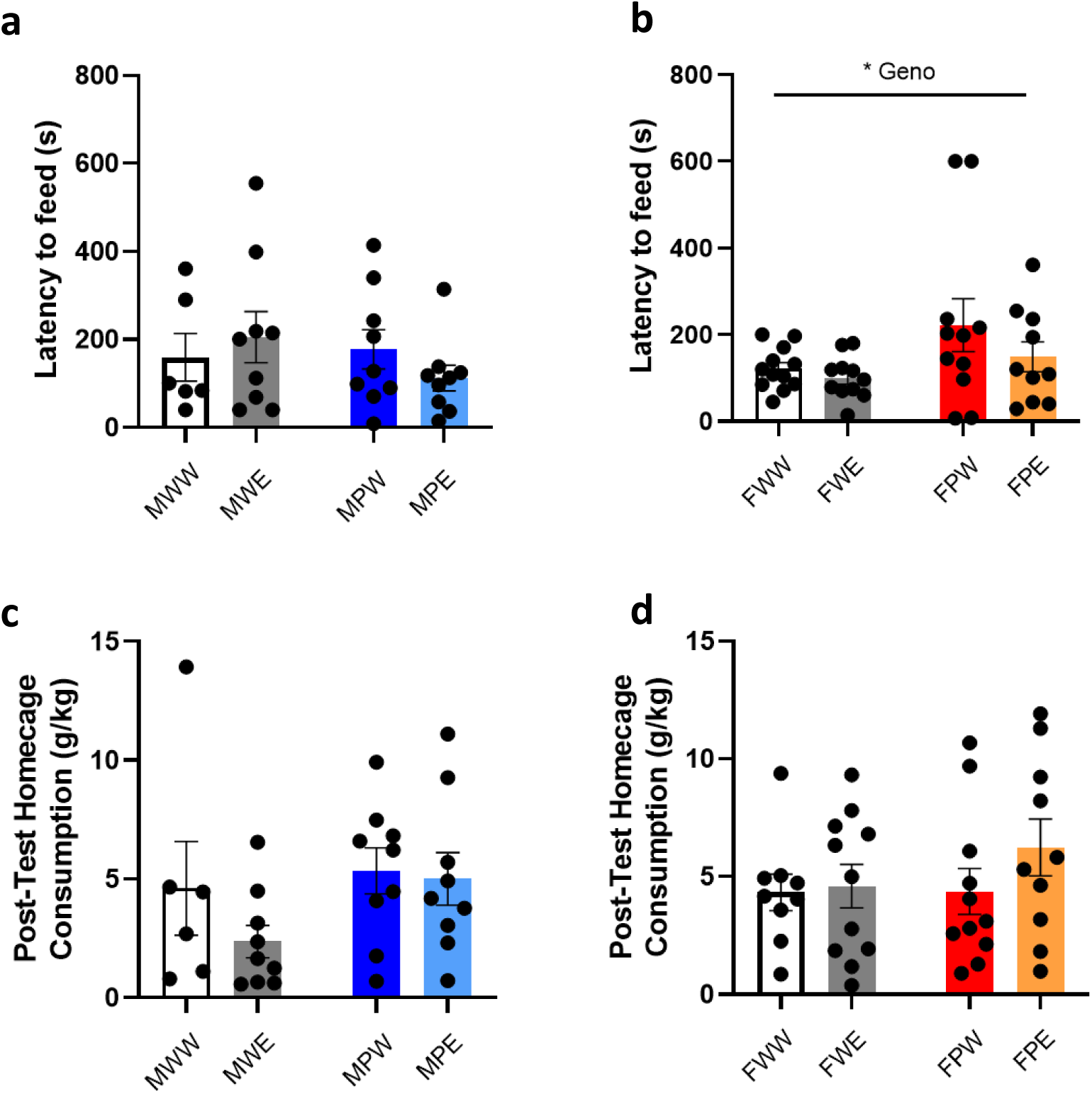
Novelty Induced Suppression of Feeding Test. **(a, b)** Latency to feed in novel environment (s). **(c, d)** Post-test homecage consumption (g/kg).

### Light-Dark Box

The LD test is broadly used as a measure of conflict between rodents’ innate aversion to (avoidance of) brightly lit areas, as well as the spontaneous exploratory behavior of rodents in response to mild stressors (Bourin and Hascoët, 2003). Male P301S mice were quicker to enter the light side (**Fig. 6a, Males latency to enter the light side**, two-way ANOVA: genotype x treatment: F (1, 40) = 0.1123, p = 0.7393; treatment F (1,40) = 0.2321, p = 0.6326 ns ; genotype F (1, 40) = 8.8828, p = 0.0050**), and spent more time in the light (**Fig. 6b, Males total duration in light side**, two-way ANOVA: genotype x treatment: F (1, 40) = 0.4624, p = 0.5004; treatment F (1,40) = 0.1308, p = 0.7195 ns; genotype F (1, 40) = 7.123, p = 0.0109*). Post-hoc Bonferroni test reveals significant differences in MWE vs. MPE for latency to enter and time in light side (**Fig. 6a**, WT-P301S, water p = 0.13825, ethanol p = 0.0495; **Fig. 6b**, Bonferroni, WT-P301S, water p = 0.3330, ethanol p = 0.0461*). Previous studies utilizing the light-dark box test have found the first 5 minutes to be the most aversive, therefore we also examined time spent in the light in 5-minute bins (Mozhui et al., 2010). Analyzing the duration in light in 5-minute bins revealed a main effect of genotype in males (**Fig. 6c Males Duration in light side, 5-min bins**, three-way ANOVA: time: F (2,80) = 6.184, p = 0.0032**; treatment F (1,40) = 0.1196, p = 0.7313; genotype F (1, 40) = 9.209, p = 0.0042**; genotype x treatment: F (1, 40) = 9.209, p = 0.4616; time x genotype x treatment: F (2,80) = 0.02959*, p = 0.9709). Post-hoc Tukey test did not reveal significant differences among male groups.

**Figure 6.**
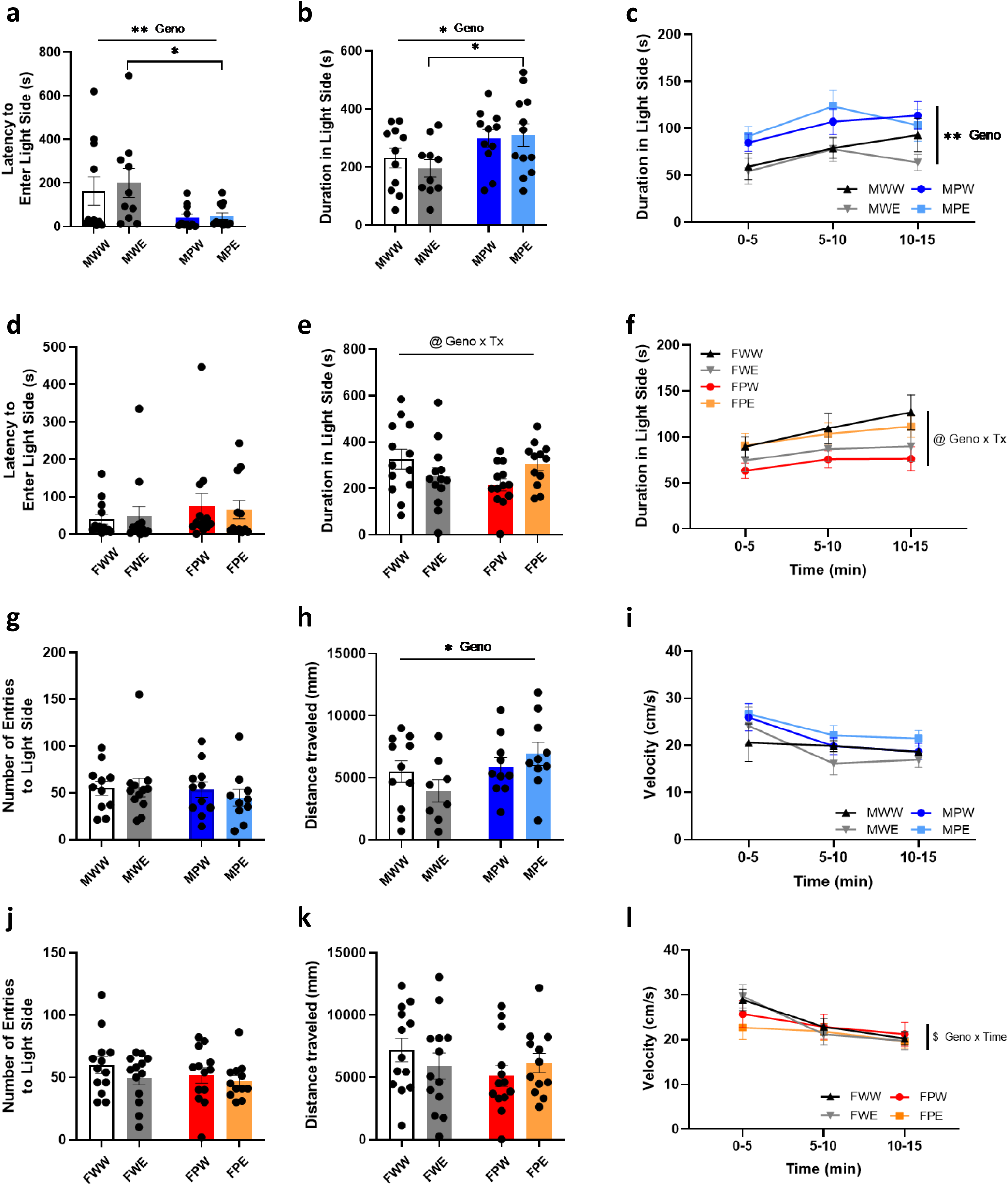
Light Dark Box. **(a)** Males latency to enter light side (s). **(b)** Males total duration in light side (s). **(c)** Males five minute binned duration in light side (s) **(d)** Females latency to enter light side (s) **(e)** Females total duration in light side (s) **(f)** Females five minute binned duration in light side (s). **(g)** Males number of entries to light side. **(h)** Males distance traveled (cm?) **(i)** Males velocity (cm/s). **(j)** Females number of entries to light side. **(k)** Females distance traveled (cm??) **(l)** Females velocity (cm/s)

In female mice, there were no significant differences in latency to enter (**Fig. 6d Females latency to enter the light side**, two-way ANOVA: genotype x treatment: F (1, 47) = 0.1358, p = 0.7141; treatment F (1,47) = 0.001303, p = 0.9714; genotype F (1, 47) = 1.089, p = 0.3020), however, there was a significant genotype by treatment interaction on time in light side when examining total time in light (**Fig. 6e Female Duration in light side**, two-way ANOVA: genotype x treatment: F (1, 47) = 5.562, p = 0.0226*; treatment F (1,47) = 0.04762, p = 0.8282; genotype F (1, 47) = 0.6436, p = 0.4264), as well as duration in light side in five minute time bins (**Fig. 6f Females Duration in light side, 5-min bins**, three-way ANOVA: time: F (2,94) = 6.992; p = 0.0015**; treatment F (1,47) = 0.04954, p = 0.8248; genotype F (1, 47) = 0.6354, p = 0.4294; genotype x treatment: F (1, 47) = 5.578, p = 0.0224*; time x genotype x treatment: F (2,94) = 0.8688; p = 0.4228). Post-hoc Tukey test of total time in light as well as duration in light side in five-minute bins did not reveal significant differences among groups.

In both male and female groups there were no significant differences in the number of transitions into the light side. (**Fig. 6g Males number of entries to light side**, two-way ANOVA, treatment: F (1,40) = 0.2319, p = 0.6327 ; genotype: F (1, 40) = 0.5240, p = 0.4733; genotype x treatment: F (1, 40) = 0.2891, p = 0.5938; **Fig. 6j Females number of entries to light side**, two-way ANOVA:; treatment: F (1,47) = 1.552, p = 0.2191 ns; genotype: F (1, 47) = 0.7905, p = 0.3785; genotype x treatment: F (1, 47) = 0.2744, p = 0.6029).

Given the observed increase in locomotor activity in the open field test, we also examined the distance traveled in the light side of the light-dark box, as well as velocity. Male P301S mice were more active in the light side as measured by distance traveled, and there was a genotype by time interaction in velocity traveled among females (**Fig. 6h Males distance traveled**, two-way ANOVA, treatment: F (1,40) = 0.2866, p = 0.5953; genotype: F (1, 40) = 5.761, p = 0.0211*; Bonferroni multiple comparison testing: WT-P301S: water t = 0.680, p > 0.9999; ethanol t = 2.710, p = 0.0197*; **Fig. 6k Females distance traveled**, two-way ANOVA, treatment: F (1,47) = 0.02229, p = 0.8820; genotype: F (1, 47) = 0.9889, p = 0.3251; genotype x treatment F (1,47) = 1.559, p = 0.2180; **Fig. 6i Males velocity in 5-min bins**, males, three-way ANOVA, time: F (2,78) = 13.42, p < 0.0001; treatment: F (1,39) = 0.08954, p = 0.7663; genotype: F (1, 39) = 2.476, p = 0.1237; **Fig. 6l Females velocity in 5-min bins**, three-way ANOVA, time: F (2,94) = 24.35, p < 0.0001; treatment: F (1,47) = 0.3257, p = 0.5709; genotype: F (1, 47) = 0.5004, p = 0.4828; post-hoc Tukey multiple comparisons: ns).

### Social Behavior

Patients with dementia and AD exhibit social withdrawal in addition to altered social behavior, sometimes referred to as disinhibition (Gottesman and Stern, 2019; Altomari et al., 2022). A previous study using a mouse line with the P301S mutation under a different promoter (called the TAU58/2 line) examined social behavior in four-month-old male mice and found reduced sociability in the social preference test, as well as reduced socio-positive behaviors such as nosing and anogenital sniffing (Watt et al., 2020). In the present study, we found no significant differences in total time interacting during the free social interaction test for male mice as well as females, however examining the data by cumulative time interacting revealed a significant treatment effect in females (**Fig. 7a, Males total time interacting**, two-way ANOVA: genotype x treatment: F (1,36) = 0.01762, p = 0.8951; genotype: F (1,36) = 0.4259; p = 0.5182; treatment F (1,36) = 0.6717, p = 0.4179. **Fig. 7b, Males cumulative time interacting**, three-way ANOVA: time x treatment: F (9, 315) = 0.9383, p = 0.4919; genotype: F (1,36) = 0.7528; p = 0.3195; treatment F (1,36) = 0.4285, p = 0.5170. **Fig. 7c, Females total time interacting**, two-way ANOVA: genotype x treatment: F (1,47) = 0.149, p = 0.7010; genotype: F (1,47) = 2.511; p = 0.9839; treatment F (1,47) = 2.511, p = 0.1198. **Fig. 7d, Females cumulative time interacting**, three-way ANOVA: time x treatment: F (9, 423) = 2.238, p = 0.0189*; genotype: F (1,47) = 0.1453; p = 0.7048; treatment F (1,47) = 2.2126, p = 0.1514; post-hoc testing not significant). **Fig. 7d** shows prior ethanol experience drives increased sociability in both WT and PS19 females; however, post-hoc multiple comparison tests did not yield significant differences.

**Figure 7.**
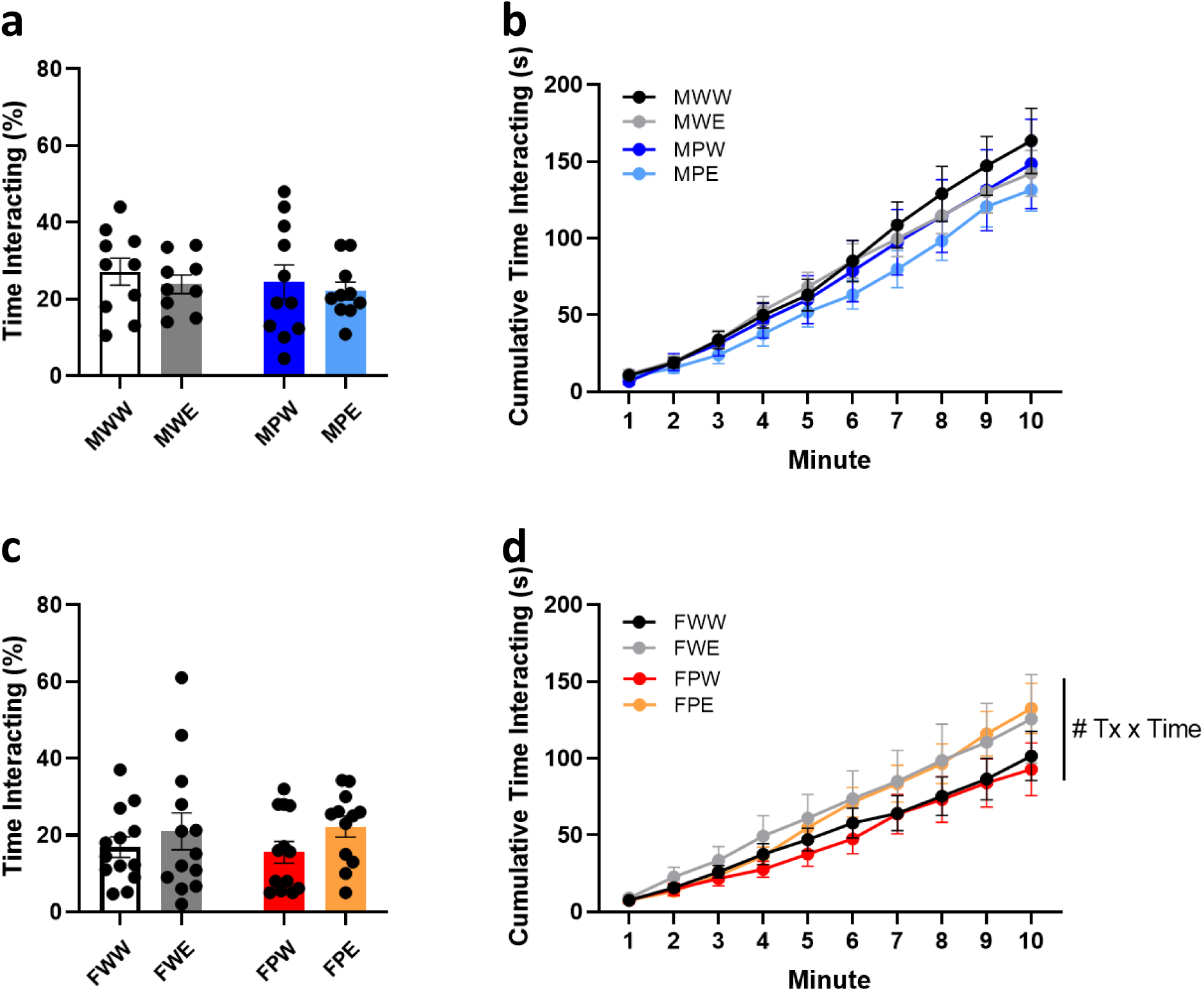
Free Social Interaction. **(a)** Males Percent Time Interacting (%). **(b)** Males Cumulative Time Interacting (s). **(c)** Females Percent Time Interacting (%). **(d)** Females Cumulative Time Interacting (s).

A closer examination of the frequency and duration of different types of prosocial and antagonistic behaviors revealed ethanol drove different effects in male and female mice, as summarized in **Table 1**.

**Table 1.**
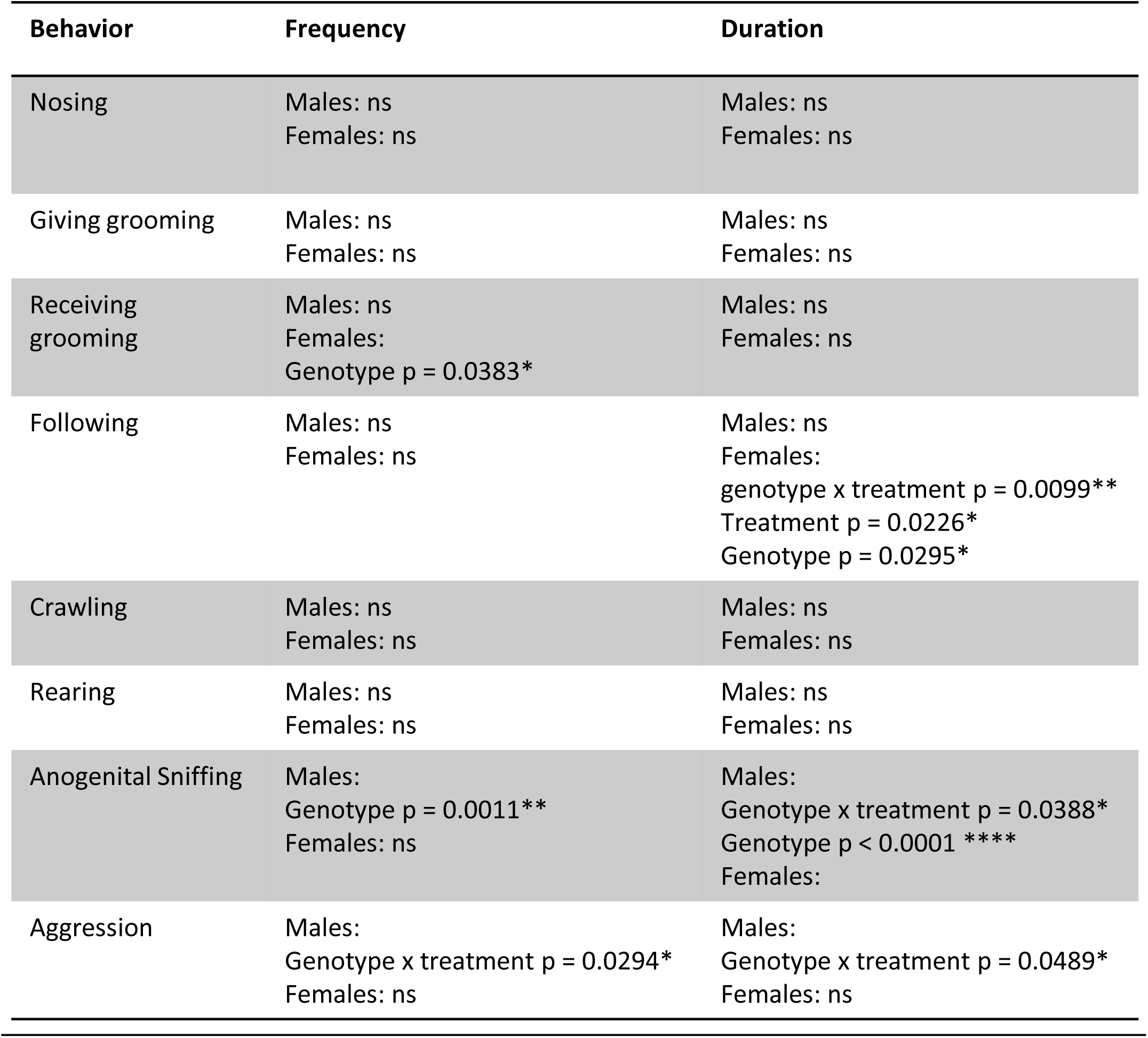
Prosocial and Antagonistic Behaviors During Free Social Interaction. For each sex, the frequency and duration of each behavior were analyzed by two-way ANOVA, followed by Bonferroni’s multiple comparisons in the event of a significant effect. P301S males exhibited increased aggression and anogenital sniffing, with ethanol amplifying this effect in P301S mice but reducing in WT. P301S females had significantly decreased frequencies of receiving grooming. P301S females also spent significantly less time following the social target mouse, with ethanol reducing this effect in P301S females but increasing this effect in WT.

Briefly, P301S males exhibited fewer periods of anogenital sniffing and ethanol enhanced aggressive behavior in P301S males, while P301S females experienced fewer periods of receiving grooming.

Male P301S mice had fewer instances and spent less time anogenital sniffing (**Table 1: Anogenital Sniffing Duration**: two-way ANOVA, genotype x treatment F (1, 36) =4.600, p = 0.0388*; genotype F (1, 36) = 19.94, p < 0.0001****; treatment F (1, 36) = 1.959, p = 0.1702. Post-hoc Tukey’s test: MPW vs. MWE p = 0.0008***; MWE vs. MPE p = 0.0002***. (**Table 1: Anogenital Sniffing Frequency**: two-way ANOVA, genotype x treatment F (1, 36) = 2.041, p = 0.1617; genotype F (1, 36) = 0.2134, p = 0.6469; treatment F (1, 36) = 12.49, p = 0.0011**. Post-hoc testing did not reveal further significant differences.)

There is a significant treatment by genotype interaction in aggression time among males (**Table 1: Aggression Duration**: two-way ANOVA, genotype x treatment F (1, 36) =4.145, p = 0.0489*; genotype F (1, 36) = 0.3562, p = 0.5544; treatment F (1, 36) = 1.551, p = 0.2210. Post-hoc testing did not reveal further significant differences).

Female P301S spent less time following the social target mouse and there was a significant genotype by treatment interaction such that ethanol exposure reduced following behavior in wildtype females (**Table 1: Following Duration:** two-way ANOVA: genotype x treatment F (1, 47) =7.222, p = 0.0099**, genotype F (1, 47) = 5.042, p = 0.0295*, treatment F (1, 47) = 5.560, p = 0.0226*. Post-hoc Tukey’s test: FWW vs. FPW p = 0.0051**, FWW vs. FWE p = 0.0041**, FWW vs. FPE p = 0.120*). Female P301S mice also had fewer instances of receiving grooming (**Table 1: Receiving Allogrooming Frequency:** two-way ANOVA: genotype x treatment F (1, 47) =7.222, p = 0.9867; genotype F (1, 47) = 4.543, p = 0.0383*; treatment F (1, 47) = 0.02688, p = 0.8705. Post-hoc testing did not reveal further significant differences).

### Learning and Memory Assays

Memory deficits have previously been reported in the P301S line, as measured by the Morris Water Maze and fear conditioning and recall (Takeuchi et al., 2011; Lasagna-Reeves et al., 2016). In the present study, novel object memory was tested in which the amount of time it took each subject to reach 20 seconds object exploration was recorded and the time spent interacting with the novel object out of those 20 seconds was reported as preference for novel object. There is a significant genotype by treatment interaction in time to reach 20-second exploration criteria such that MPW mice took longer to explore than MPE mice (**Fig. 8a, Time to reach 20s object exploration:** males: two-way ANOVA: genotype x treatment: F (1,38) = 5.546, p = 0.0238*; genotype: F (1,38) = 3.840; p = 0.0676 ; treatment F (1,38) = 0.3074, p = 0.5825). Post-hoc Bonferroni multiple comparison testing revealed a significant difference between wildtype and P301S male mice exposed to ethanol (Tukey’s: MPW vs. MPE, p = 0.0229*). When comparing between conditions, there are no deficits in preference for the novel object across any of the male groups (**Fig. 8b, Preference for novel object:** males: two-way ANOVA: genotype x treatment: F (1,38) = 1.277, p = 0.2654; genotype: F (1,38) = 0.5428 ; p = 0.4666 ; treatment F (1,38) = 2.207, p = 0.1456). When we examined if novel object preference differed from chance (one sample t-test) however, the only group that did not show a significant preference for the novel object were the MPE mice (MWW p = 0.0133* t = 3.000 df = 10, MWE p = 0.0100* t = 3.3542 df = 8, MPW p = 0.0211* t = 2.7333 df = 10, MPE p = 0.6623 t = 0.4500 df = 10.) Together, these data suggest ethanol exposure differentially affects male P301S mice, such that MPE mice explore the novel object sooner, but have reduced preference for the novel object in the novel object recognition test.

**Figure 8.**
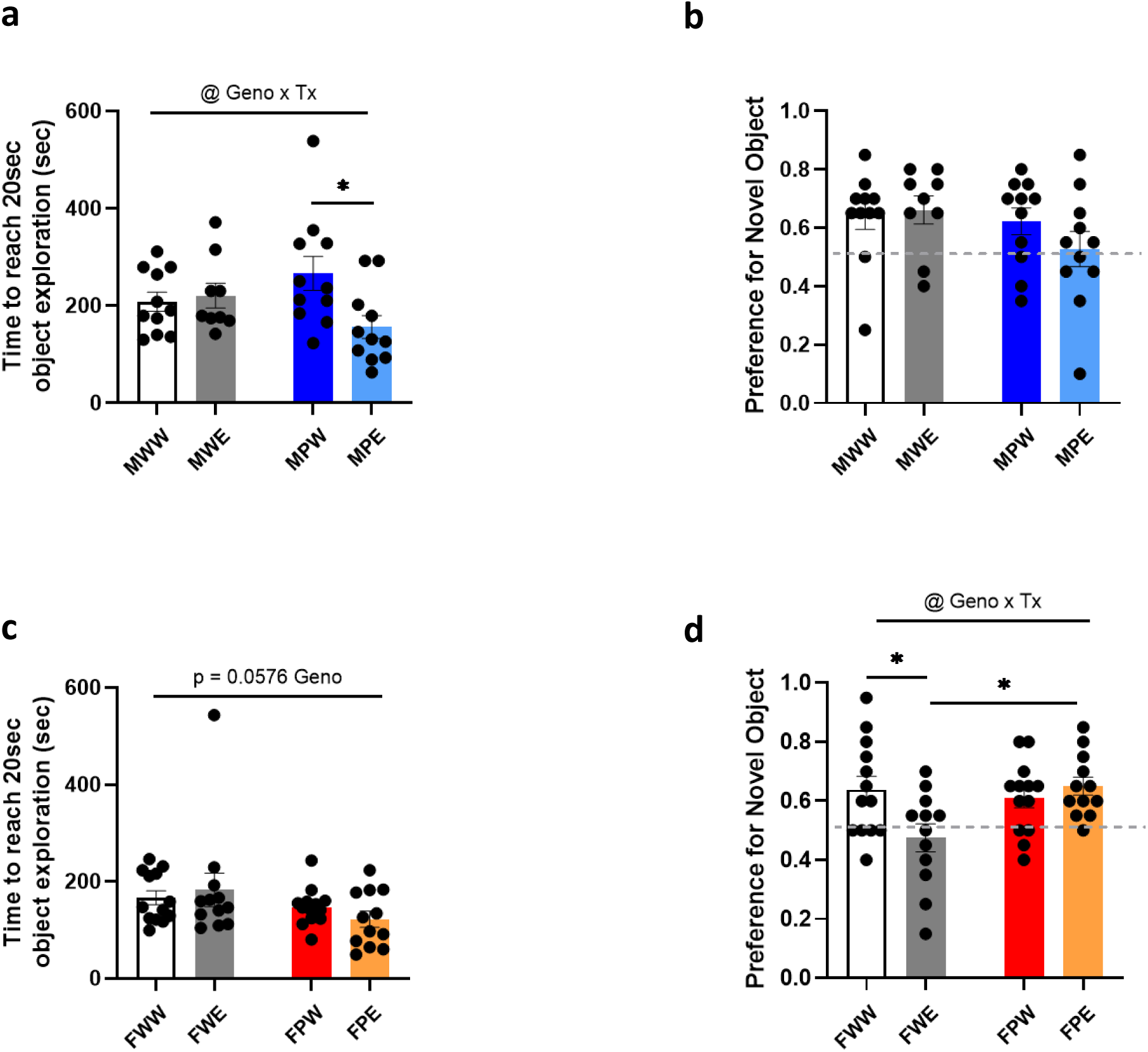
Novel Object Recognition. **(a, c)** Time to Reach 20 Seconds Object Exploration (s). **(b, d)** Males Preference for Novel Object. Dotted line (0.5) indicates equal preference for novel and familiar object. Value > 0.5 = preference for novel object. Value < 0.5 = preference for familiar object.

In female mice, there was a trend towards reduced exploratory behavior in P301S animals (**Fig. 8c, Time to reach 20 sec object exploration:** females: two-way ANOVA: genotype x treatment: F (1,46) = 1.043, p = 0.3125; genotype: F (1,46) = 0.04829; p = 0.8270; treatment F (1,46) = 3.791, p = 0.0576). There is a significant genotype by treatment interaction in novel object preference among female mice, such that FWE mice have impaired object memory while FPW and FPE do not. (**Fig. 8d, Preference for novel object:** females: two-way ANOVA: genotype x treatment: F (1,46) = 6.367, p = 0.0181*; genotype: F (1,46) = 6.367; p = 0.1251; treatment F (1,46) = 3.424, p = 0.0707). Post-hoc Tukey test revealed a significant difference between female wildtype mice exposed to water and ethanol, as well as a difference between wildtype-ethanol and P301S-ethanol mice (Tukey’s: FWW vs. FWE, p = 0.0348*; FWE vs. FPE, p = 0.0236*). We also examined if preference differed from chance in the female mice, congruent with data above female wildtype ethanol drinking mice did not show a significant preference for the novel object while all other groups did. (One sample t-test: FWW p = 0.0098** t = 3.0667 df = 12, FEW p = 0.6054 t = 0.5319 df = 11, FPW p = 0.0064** t = 3.2941 df = 12, FPE p = 0.0005** t = 4.8387 df = 11). Together this data suggest ethanol exposure differentially affects female wildtype and P301S mice.

### Learning

Next the mice underwent fear conditioning followed by context-dependent one day post-conditioning and cue-dependent recall tests two days post-conditioning. Impaired context-dependent recall in male P301S mice has previously been reported (Lasagna-Reeves et al., 2016). Male P301S spent significantly less time freezing during fear conditioning across the three conditioned stimuli, and there was a significant genotype by time interaction (**Fig. 9a. Fear learning: time spent freezing (%), males:** three-way ANOVA: genotype F (1,36) = 3.900, p = 0.0560; treatment F (1,26) = 0.0068, p = 0.9347; time x genotype: F (2,72) = 4.834., p = 0. 0107*; time x treatment x genotype F (2,72) = 2.657, p = 0.0770). However, post hoc testing (Tukey’s) did not reveal significant differences in freezing at each of the three conditioned stimuli. Together these data suggest impaired fear learning in male P301S mice. There were no significant differences in fear learning among female mice (**Fig. 9b. Fear learning: time spent freezing (%), females:** three-way ANOVA: genotype F (1,36) = 0.3187, p = 0.5759; treatment F (1,36) = 0.4699, p = 0.4974; time x genotype: F (2,72) = 1.247., p = 0. 2934; time x treatment x genotype F (2,72) = 0.6131, p = 0.5445).

**Figure 9.**
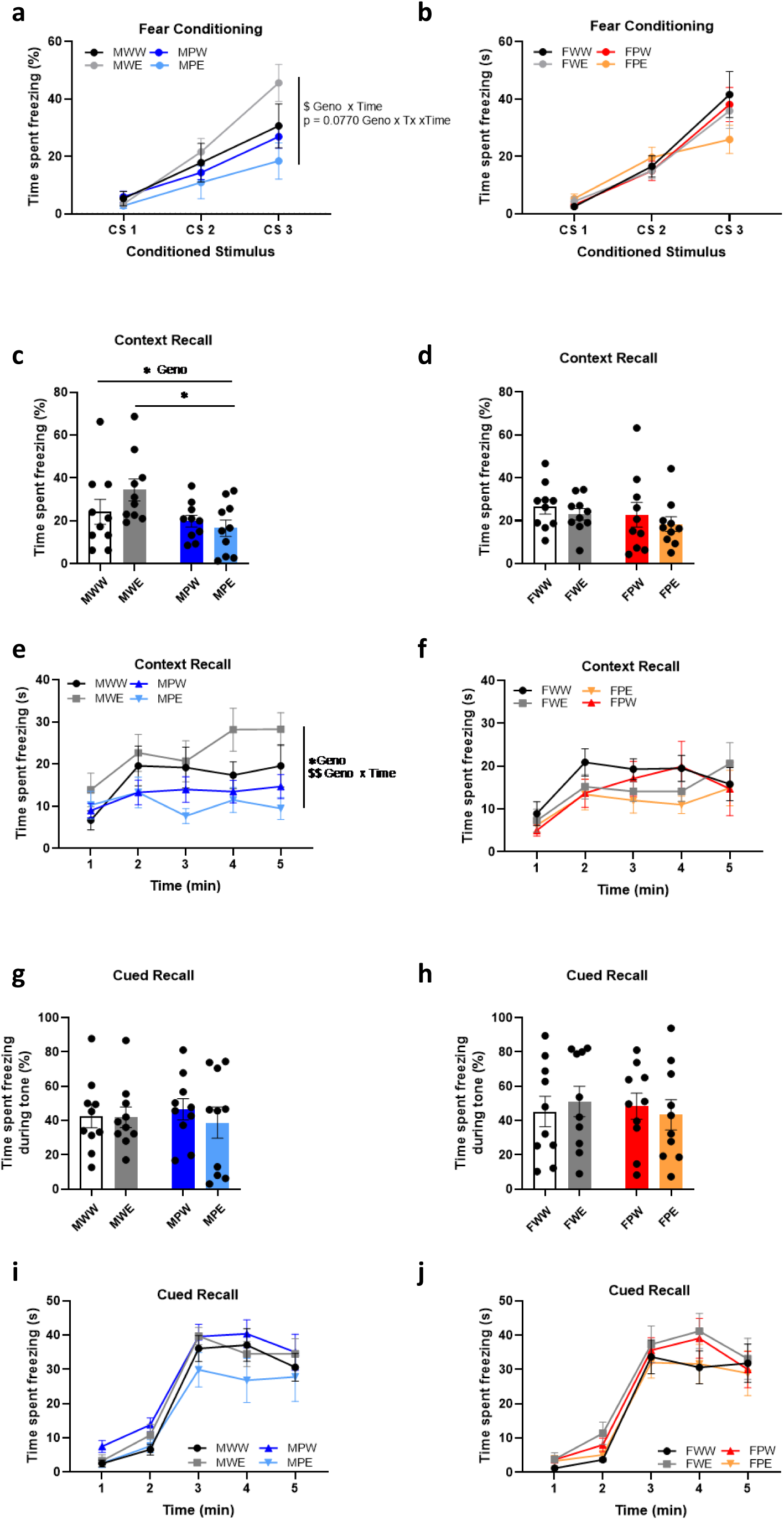
Fear Learning. **(a, b) Fear Learning**: time spent freezing during conditioned stimulus (%). **(c, d) Context-dependent Recall:** total time spent freezing (%). **(e, f) Context-dependent Recall by Minute:** time spent freezing (s). **(g, h) Cue-dependent Recall:** total time spent freezing (%). -way ANOVA, ns. **(i) Cue-dependent Recall by Minute:** time spent freezing (s).

### Context-dependent recall

Given the impairment in fear learning among male P301S mice, we expected to see impaired context-dependent recall. There were significant genotype effects on total time freezing during the context recall test for males (**Fig. 9c. Context dependent recall, males:** two-way ANOVA: genotype x treatment F (1,36) = 2.217, p = 0.1452; treatment F (1,36) = 0.5945, p = 0.4457; genotype F (1,36) = 6.042, p= 0.0189*) but not females (two-way ANOVA: interaction F (1,36) = 0.01288, p = 0.9103; treatment F (1,36) = 0.9683, p = 0.3317; genotype F (1,36) = 1.073, p= 0.3072). Post-hoc multiple comparison analysis found significant differences between WT Ethanol and P301S Ethanol male mice (Bonferroni, WT-P301S, water p = 0.9951; ethanol p = 0.0167*). There were no significant differences found in post-hoc analysis (Bonferroni) of female context recall data (**Fig. 9d. Context dependent recall, females:** two-way ANOVA, ns). Analysis of freezing over time during context-recall revealed significant differences in male wildtype and P301S mice (**Fig. 9e. Context dependent recall by minute, males:** three-way ANOVA: genotype F (1,36) = 6.935, p = 0.0124*; treatment F (1,36) = 0.4003, p = 0.5309; time x genotype: F (4,144) = 3.690, p = 0. 0068**). Post-hoc analysis (Tukey’s) did not reveal significant effects at any given time point. In female mice, analysis of freezing over time during context-recall did not identify any significant differences due to genotype and/or treatment (**Fig. 9f. Context dependent recall by minute, females:** three-way ANOVA, ns). Together this data suggest male P301S mice have impaired context-dependent memory related to impaired fear learning compared to wildtype, while female P301S and wildtype mice do not have deficits in context-dependent recall.

### Cue-dependent recall

There were no significant differences in total time freezing during the first cued recall test in males (**Fig. 9g. Cue dependent recall, males:** two-way ANOVA: genotype x treatment F (1,36) = 0.2501, p = 0.6200; treatment F (1,36) = 0.3391, p = 0.5640; genotype F (1,36) = 0. 0040, p= 0.9497) or females (**Fig. 9g. Cue dependent recall, males:** two-way ANOVA: genotype x treatment F (1,36) = 0.4012, p = 0.5304; treatment F (1,36) = 0.003033, p = 0.9564; genotype F (1,36) = 0.07875, p= 0.7806). Analysis of freezing over time during cued-recall did not identify any significant differences due to genotype and/or treatment in males nor females (**Fig. 9i, j. Cue dependent recall by minute)**. Together these data suggest intact cue-dependent fear learning.

## Discussion

In this study we examined drinking behavior, locomotor activity, multiple aspects of anxiety-like behavior, social behavior, and learning and memory in male and female P301S mice and wildtype littermate controls following 16 weeks of intermittent access to ethanol. We identified significant differences across the domains of sex, genotype, and treatment (ethanol consumption). Notably, female mice of both genotypes drank significantly more than male mice, however in assays where we identified significant genotype-treatment interactions among females, P301S ethanol-naïve mice (FPW) exhibit behavioral deficits, while ethanol consumption among P301S females (FPE) appears to result in behavioral phenotypes similar to control mice (FWW) and wildtype-ethanol mice (FWE). We propose these findings lay the groundwork to further examine the interaction between ethanol consumption and tauopathy.

### Difference in drinking behaviors in male and female mice, yet similar consumption across genotypes

Examination of drinking behavior under the IA paradigm revealed no differences in total ethanol consumption between each genotype within each sex, however differences in total fluid intake among both P301S mice of both sexes, water intake among wildtype and P301S females, and preference for ethanol in both sexes of P301S mice suggest altered drinking behavior compared to wildtype. Female mice of each genotype drank significantly more ethanol than males of each genotype, however the difference in male and female drinking aligns with previous studies in C57B/6J mice where female mice have been observed to drink more than males (Hwa et al., 2011). Further examination of drinking on a weekly basis revealed an interaction between genotype and week in both average daily ethanol consumption per week and cumulative ethanol consumption per week, which suggests that as time went on, male P301S mice consumed more ethanol than their WT littermates. Although post-hoc testing did not identify further significant differences between the male genotypes across individual weeks, extending drinking periods beyond 16 weeks may reveal greater consumption in the male P301S animals. This hypothesis is also supported by a significant genotype by week interaction in fluid intake but no differences in total water intake. The female data suggests when examining preference for ethanol bottle, there is a significant main effect of genotype, such that female P301S mice have greater preference for the ethanol bottle, driven by reduced water intake. While post-hoc testing did not reveal significant differences between the female genotypes on a week-to-week basis, extended drinking periods may have shown increased ethanol preference in the female P301S animals.

#### Limitations of Intermittent Access Paradigm

IA requires the animals to voluntarily consume ethanol, which may limit dosage, and limits comparisons between groups that drink different quantities (i.e. male and female mice). Further studies could use experimenter administered models (such as administration by injection, inhalation, or oral gavage) to understand phenotypic responses in a dose and blood alcohol concentration-dependent manner. This would allow for the exploration of alcohol dependence as well within the P301S mice. While the IA paradigm enables voluntary consumption of ethanol, and we found changes in both male and female mice in terms of patterns of consumption and preference for ethanol, it does not assess the subject’s motivation to consume ethanol. Future studies could also examine operant ethanol self-administration with fixed and progressive ratios to determine whether the P301S line i exhibits enhanced motivation compared to littermate wildtype controls. Such studies could examine self-administration with or without prior history of ethanol exposure, as well as examine self-administration across the lifespan. These studies may reveal insights as to whether the timeline of pathological changes in this mouse line correlate with changes in ethanol self-administration, mirroring what we are observing with our preference data in male mice. Interestingly, a paper describing a mouse line expressing alpha synuclein (αSYN), a major genetic contributor to Parkinson’s Disease, reported no differences in ethanol consumption or preference compared to wildtype, however operant conditioning experiments revealed the transgenic αSYN mouse line showed higher motivation for ethanol and attenuated context and cued-induced reinstatement of alcohol-seeking behavior (Rotermund et al., 2017). Their data suggest the progression of Parkinson’s Disease-like pathology is associated with the primary reinforcing effects of ethanol, which is interesting given overlap in brain regions affected by both Parkinson’s Disease and Alzheimer’s Disease (Butkovich et al., 2020). Future studies of alcohol consumption in neurodegenerative mouse models may elucidate directionality: does ethanol contribute to neurodegeneration, and does neurodegeneration drive ethanol consumption?

### Effects of Ethanol and Tauopathy in Males

Although male mice consumed significantly less ethanol than female mice in this study, statistical analyses within male groups revealed significant genotype by treatment interactions. Interestingly, the only significant main effect of ethanol experience in male mice was in locomotor activity in the open field test (Fig 2b, c). However, across several measurements in males there were significant main effects of genotype: P301S male mice (MPW and MPE) had ***attenuated growth curves*** over 16 weeks of IA than wildtype controls (MWW and MWE) (Fig. 2a); were ***more active*** in the open field test (Fig 3a, b, c) and light-dark box (Fig. 6h); ***consumed differential amounts of ethanol and had differential ethanol preference over 16 weeks of IA (but no differences in total consumption)*** (Fig. 2d, e, f); exhibited ***reduced avoidance behavior*** in the EPM (Fig. 4a, c) and light-dark box (Fig. 6a, b, c); behaved ***less prosocial and more antagonistic*** during free social interaction (Table 1); and presented ***impaired fear learning and context-dependent recall*** (Fig. 9a, c, e). Across several measures, there were significant genotype by treatment interactions in which ethanol had opposing effects in wildtype and P301S males. Notably, ethanol significantly exacerbated some phenotypical differences in the MPE group, such as ***hyperactivity*** in the open field (Fig. 2a, b), ***quicker exploration*** in the novel object memory task (possibly due to increased hyperactivity) (Fig. 9a), and ***increased prosocial (anogenital sniffing) and antagonistic (aggressive) behavior*** (Table 1). Hyperactivity in approach/avoidance assays and mazes often is a confounding variable in data interpretation. Interestingly, while it has been previously reported male P301S mice are hyperactive at four months of age (Takeuchi et al., 2011), the hyperactivity we observed in our study only occurred in contexts where there was bright lighting (40 lux +). When we observed mice in low lighting conditions (9.5 lux) in the EPM, we did not see differences in distance traveled or closed arm entries. Our data suggests that the “hyperactivity” observed in P301S mice is dependent on environmental stressors. Ethanol consumption decreased the time it took P301S mice to explore the novel object, however these mice did not show preference for the novel object which suggests that ethanol consumption conferred a memory deficit in P301S mice.

Together these data suggest tauopathy conferred by the P301S transgene in male mice results in several behavioral deficits, further altered by ethanol consumption. Based on our findings, we hypothesize increased dosing of ethanol such as by using an involuntary consumption model, or increasing the time course of ethanol consumption and examining mice at a later age may yield more genotype-treatment interactions.

### Effects of Ethanol and Tauopathy in Females

Statistical analyses within female groups revealed significant main effects of genotype, treatmen, and genotype by treatment interactions. Across several measurements in females there were significant main effects of treatment as well as treatment by time interactions: Female ethanol drinkers (FWE and FPE) exhibited ***more weight gain*** over 16 weeks of IA than wildtype controls (FWW and FPW) (Fig. 2b); and spent ***more time socializing*** in the free social interaction test (Fig. 7d).

Across several measurements in females there were significant main effects of genotype: P301S female mice had ***increased preference for the ethanol bottle*** and ***reduced total fluid and water intake*** compared to wildtype (Fig. 2f, g-j) although no difference in total, weekly, or cumulative ethanol consumption (Fig. 2c, d, e); ***differences in approach/avoidance behavior across different contexts***, including a trend towards reduced avoidance behavior in the EPM (Fig. 3f) but increased avoidance in the NSF test (Fig. 5b); ***reduced prosocial behaviors*** (Table 1); and ***impaired cue-dependent recall*** following fear conditioning (Fig. 9h).

Given that female mice consumed more ethanol than males, we predicted more instances of genotype by treatment interactions across behavioral measures in female mice. Surprisingly, the significant genotype by treatment interactions seen among female mice seemed to confer more “resiliency” due to P301S genotype in behavioral deficits than what we observed in males. In the following measures we observed the combination of tauopathy and ethanol consumption in females (FPE) resulted in behavioral phenotypes mores similar to controls (FWW), while inducing alterations in the ethanol treatment group (FWE) and the P301S group (FPW): Notably, in the open field test, ***FPE mice habituate to the arena more so than FPW mice*** (Fig. 3e, f); ethanol consumption differentially affected avoidance behavior in the LD box test, driving increased avoidance behavior in wildtype (FWE) but ***reduced avoidance behavior in P301S*** (FPE) (Fig. 6e, f); ethanol consumption also differentially affected novel object memory, driving reduced preference for the novel object (impaired memory) in FWE mice but seemingly ***improved memory in FPE*** (Fig. 8d, genotype x treatment p = 0.0151*).

### Considering Ethanol Consumption, Severity in Tauopathy, and Sex as Variables on the Biological Spectrum

There are a few possible interpretations of these data when considering both males and females. In instances where there is a main effect of genotype in males but genotype by treatment interaction in females, there are at least three plausible explanations 1) <u>differences in ethanol consumption</u>: perhaps there is a dose-dependent threshold at which ethanol consumption impacts the behaviors assessed in this study, and the male mice did not consume enough ethanol to reach this threshold, 2) <u>differences in tauopathy:</u> the severity of tauopathy may differ in male and female mice at the timepoint at which we performed our studies, and 3) <u>sex as a biological variable:</u> differences in male and female biology may contribute to the differential effects of tauopathy and ethanol on behavior observed in this study. Future studies can use a non-contingent model of ethanol consumption in order to control ethanol dose, BACs, and better assess sex as a biological variable. Future studies can also assess tau levels in individual mice to account for differences in severity of tauopathy. Since this goal of this study was to examine how long-term, voluntary ethanol consumption affects behavior in wildtype and P301S mice of both sexes, we examined the results separately for each sex, and in the context of BPSD.

### Implications for the HIDA Domain of the BPSD

The HIDA, or HIIDAA, domain of BPSD includes <u>h</u>yperactivity, impulsivity, irritability, <u>d</u>isinhibition, <u>a</u>ggression, and <u>a</u>gitation (Keszycki et al., 2019). In patients, symptoms may present as inappropriate inhibition of motor responses resulting in wandering, or inappropriate inhibition of thoughts resulting in inappropriate speech and social behavior (reviewed in Keszycki et al. 2019). Several behavioral assays in the present study address these features. Activity levels were assessed in the open field, EPM, and LD box tests; aggression was assessed in the free social interaction test; and disinhibition was assessed in assays broadly used to measure approach/avoidance behavior including the EPM, LD box, and NSF test.

#### Hyperactivity

The P301S mouse line has previously been characterized in terms of locomotor activity, with mixed reports of differences in activity compared to wildtype littermate controls. Two studies noted hyperactivity in males beginning as early as 2 months of age (Scattoni et al., 2010; Takeuchi et al., 2011). One study noted no differences in locomotor activity until 5 months of age, where they found P301S mice were less active than WT (Xu et al., 2014).

In this present study, locomotor hyperactivity was observed in the open field test and LD box, but not the EPM, although hyperactivity exhibited in different manifestations in male (increased total distance traveled) and female tauopathy mice (increased velocity). It is possible hyperactivity was noted in the open field and LD box but not the EPM due to differences in light level in the assays (an external stressor in the environment)-the OF and LD box are brightly lit (≥40 lux) whereas the EPM is not (9.5 lux).

#### Disinhibition

In this study we observed disinhibition (an inability to inhibit behaviors in a given setting or circumstance) in both social and non-social settings. In the free social interaction test, ethanol exposure increased sociability in wildtype and P301S females. In both male and female P301S mice, we observed altered social behavior types. In non-social settings, male P301S mice (both MPW and MPE) presented increased disinhibitory behavior in the EPM and LD box, while female P301S mice had increased inhibition in the NSF test and LD box, and the FPE group had reduced inhibition (increased disinhibition) in the LD.

#### Aggression

Several transgenic tauopathy and amyloidosis mouse models exhibit aggression (reviewed in Kosel et al., 2020). While we did note increased aggression among the MPE group in the free social interaction test, and removed mice from the test who fought for over ten seconds, the resident-intruder task is considered the standard for assessing aggression in mice and would be better suited to draw conclusions (Kosel et al., 2020).

#### Agitation

Early rodent studies examined hyperactivity through the lens of agitation and restlessness (Lister, 1987; Carola et al., 2002). A review by Kosel et al., of several transgenic tau and amyloidogenic mouse models found mixed results in terms of agitation assessed during anxiety-like behavior assays (such as open field, EPM, and LD box) and locomotor activity (which also includes open field testing, as well as home-cage testing). The authors of the review note it is difficult to assess agitation in mice since agitation is likely related to avoidance behavior and exploratory behavior. In the present study, increased locomotion was only observed in brightly lit environments but not the EPM. Especially because the greatest differences in locomotion are noted in the first binned time epoch of the OF (10 min) and LD (5 min), but no differences in locomotion were noted in the 5-min EPM test, our data suggest increased agitation in brightly lit, aversive environments.

### Implications for Anxiety and Fear, Two Domains of the BPSD

#### Anxiety

By using multiple behavioral assays, we were able to sample the nuance of anxiety-like behavior in mice, including approach/avoidance and inhibition/disinhibition in environments considered aversive to rodents: novel, elevated, open and/or brightly lit spaces. The free social interaction test has also previously been used (and pharmacologically validated) to assess anxiety, based on increases in sniffing, following, grooming, and fighting (boxing and wrestling) (Pentkowski, File and Hyde 1978, File and Seth 2003).

Male P301S mice exhibited what has classically been interpreted as reduced anxiety-like behavior across several tests, EPM and LD, as well as more non-traditional tests such the free social interaction test. The reduced anogenital sniffing in male P301S mice may also be interpreted as reduced anxiety-like behavior in the presence of a novel mouse. We and others, however, have observed similar changes in behavior following stress and in withdrawal from drugs in C57BL/6J mice, which has led us to interpret these changes as behavioral disinhibition (Osborn et al., 1998; Lacroix et al., 2000; Mozhui et al., 2010; Masneuf et al., 2014; Fish et al., 2018; Lowery-Gionta et al., 2018; Bloch et al., 2020; Bravo et al., 2020) Interestingly we did not see a significant main effect of treatment nor genotype by treatment interaction in male mice across these assays. It is possible the IA paradigm is insufficient to dysregulate negative affective behavior. A previous study in C57BL/6J mice following seven weeks of IA reported mild disinhibition in the EPM, but no significant differences between ethanol-drinking and ethanol-naïve males in the NSF test (Bloch et al., 2020). It is likely that if the P301S mice were on a different genetic background that there may be different results in these assays, however we used mice backcrossed to the C57BL/6J strain because of their propensity to consume ethanol, and we specifically wanted to observe how freely consumed ethanol could alter these behavioral phenotypes.

Among female P301S mice, there were several observations of increased anxiety-like behavior, and in some cases ethanol consumption reducing anxiety-like behavior. Interestingly, the significant interaction between genotype and treatment was seen in tests with brightly lit environments (open field and LD box), while a significant main effect of genotype was seen in the NSF test, and there were no significant differences in the EPM, in which both the NSF and EPM arenas are more dimly lit environments (≤10.5 lux). Notably, in the open field test, FPW mice did not habituate to the arena, unlike the FWW, FWE, and FPE groups (**Fig. 3e, f**). Ethanol consumption differentially affected anxiety-like behavior in the LD box test, driving increased anxiety-like behavior in the FWE group but reduced anxiety-like behavior in FPE group (**Fig. 6e, f**). In contrast, in the NSF test, both the FPW and FPE groups exhibited increased latency to feed compared to wildtype groups.

Furthermore, in the free social interaction test, ethanol consumption in both WT and P301S females resulted in increased cumulative time spent socializing, which may be interpreted as increased approach behavior as reduced anxiety-like behavior. When assessing behavior types by duration, FPW and FPE mice spent less time following the target conspecific mouse compared to wildtype groups, also indicating reduced anxiety-like behavior, and there was a significant genotype by treatment interaction although post-hoc multiple comparison analysis only revealed a significant difference between the FWW vs. FWE groups and not FPW vs. FPE. When examining behavior types by frequency, the FPW and FPE groups had significantly fewer instances of receiving grooming from the target conspecific mice, but no differences in performing grooming; increased grooming typically indicates increased anxiety-like behavior (Hart et al., 2010). Overall, the female data from the free social interaction test, when interpreted in the context of anxiety-like behavior, are difficult to draw a unifying conclusion from. Future studies may gain insights from investigating female mice that are not singly housed, cohabitating/interacting with cage mates in a homecage environment vs. a distinct chamber. Together these data suggest history of ethanol consumption in P301S females may result in more resiliency to brightly-lit, open space, aversive environments, resulting in reduced anxiety-like behavior.

### Implications for Social Behavior in Dementia

One of the earliest signs of dementia is social withdrawal. In some literature, social withdrawal is also considered a form of apathy. The aforementioned review by Kosel et al. noted several mouse models of AD do not exhibit general apathy, but do show increased social withdrawal across home-cage testing, free social interaction test, and the three-chamber test, and that in some models sociability is further reduced with age and progressive severity of AD-like pathology.

In our study, there were significant genotype by treatment interactions in duration of anogenital sniffing and aggressive behavior (excluding mice that were removed after ten consecutive seconds of fighting) (**Table 1**). Notably, male P301S mice spent less time anogenital sniffing compared to WT, and ethanol increased sniffing among MWE but reduced sniffing among MPE groups. Similarly, ethanol consumption had differential effects in aggressive behavior by genotype, such that MPE mice were more aggressive (although while there was a significant interaction, there was no main effect of genotype or treatment and no significant results in post-hoc multiple comparison tests, which may have been due to decreased power from removing overly aggressive animals). Together these data suggest tauopathy and ethanol consumption may alter social behavior in male mice. Female ethanol drinkers (FWE and FPE) spent more time socializing in the free social interaction test (**Fig. 7d)**. Female P301S mice (FPW and FPE) exhibited reduced prosocial behaviors (**Table 1**), including reduced time spent following the target conspecific mouse and reduced frequency of receiving grooming. We suggest further studies utilizing other types of tests to better examine social behavior, such as the three-chamber test and resident-intruder test may yield valuable insights. Additionally, by nature of the IA paradigm, our animals were singly housed for most of their lives. It would be intriguing to observe if sociability assays with novel target mice would be different depending on cohousing with conspecifics.

### Implications for Learning and Memory in Dementia

In the present study we found ethanol consumption induced impairment in object memory in P301S male mice and female WT mice, and impaired fear learning and context-dependent recall in P301S males. These data suggest that the interaction between ethanol and tauopathy is complicated, and may differ in male and female animals. However, the data especially in the males in the novel object test, does have a degree of variance, which may imply individual variability and vulnerability that should be further explored. An important consideration in our studies is that the P301S mice survival curve exhibits 50% mortality at 9 months of age (Yoshiyama et al., 2007), a time estimated to correspond to “early middle age” in humans (Flurkey et al., 2007). This restricted our ability to investigate longer ethanol consumption paradigms and how tauopathy affects “old age” in this model. We suggest the timepoint at which we performed behavior testing, however, may represent a key period in understanding BPSD before dramatic learning and memory impairment, however further cognitive testing is warranted.

Together the data presented in this study support the need for understanding the ways in which substance use and dementia intersect to influence a variety of behaviors. With an aging population and rates of alcohol use and dementia both on the rise, it is imperative for future work to explore the ways substance use engages neural circuitry involved in the behavioral and psychological symptoms of dementia, in addition to the cognitive symptoms. The onset and progression of BPSD represent a potential timepoint for intervention to treat both neuropsychiatric and neurological symptoms, and we propose understanding the motivation underlying substance use in the elderly may reveal new opportunities in screening for BPSD and early cognitive deficits associated with dementia.

## ACKNOWLEDGEMENTS

This work was supported by U01AA020911-08S1 (ZAM), the University of North Carolina Dissertation Completion Fellowship (CMC), and T3AA2007573 (AMD). We thank Graydon Gereau, Madigan Bedard, Ali Alvarez, Leslie Hassanein, and Sofia Neira for assistance with the drinking studies. We thank Graham Diering for providing mice and helpful discussion.

## References

Altomari N, Bruno F, Laganà V, Smirne N, Colao R, Curcio S, Di Lorenzo R, Frangipane F, Maletta R, Puccio G, Bruni AC (2022) A Comparison of Behavioral and Psychological Symptoms of Dementia (BPSD) and BPSD Sub-Syndromes in Early-Onset and Late-Onset Alzheimer’s Disease. J Alzheimers Dis 85:691–699.

Araujo I, Henriksen A, Gamsby J, Gulick D (2021) Impact of Alcohol Abuse on Susceptibility to Rare Neurodegenerative Diseases. Frontiers in Molecular Biosciences 8 Available at: https://www.frontiersin.org/article/10.3389/fmolb.2021.643273 [Accessed March 31, 2022].

Barnett A, David E, Rohlman A, Nikolova VD, Moy SS, Vetreno RP, Coleman LG (2022) Adolescent Binge Alcohol Enhances Early Alzheimer’s Disease Pathology in Adulthood Through Proinflammatory Neuroimmune Activation. Front Pharmacol 13:884170.

Bessey LJ, Walaszek A (2019) Management of Behavioral and Psychological Symptoms of Dementia. Curr Psychiatry Rep 21:66.

Bloch S, Rinker JA, Marcus MM, Mulholland PJ (2020) Absence of effects of intermittent access to alcohol on negative affective and anxiety-like behaviors in male and female C57BL/6J mice. Alcohol 88:91–99.

Bourin M, Hascoët M (2003) The mouse light/dark box test. Eur J Pharmacol 463:55–65.

Bravo IM, Luster BR, Flanigan ME, Perez PJ, Cogan ES, Schmidt KT, McElligott ZA (2020) Divergent behavioral responses in protracted opioid withdrawal in male and female C57BL/6J mice. Eur J Neurosci 51:742–754.

Breslow RA, Castle I-JP, Chen CM, Graubard BI (2017) Trends in Alcohol Consumption Among Older Americans: National Health Interview Surveys, 1997 to 2014. Alcohol Clin Exp Res 41:976–986.

Bugiani O, Murrell JR, Giaccone G, Hasegawa M, Ghigo G, Tabaton M, Morbin M, Primavera A, Carella F, Solaro C, Grisoli M, Savoiardo M, Spillantini MG, Tagliavini F, Goedert M, Ghetti B (1999) Frontotemporal dementia and corticobasal degeneration in a family with a P301S mutation in tau. J Neuropathol Exp Neurol 58:667–677.

Butkovich LM, Houser MC, Chalermpalanupap T, Porter-Stransky KA, Iannitelli AF, Boles JS, Lloyd GM, Coomes AS, Eidson LN, Rodrigues MEDS, Oliver DL, Kelly SD, Chang J, Bengoa-Vergniory N, Wade-Martins R, Giasson BI, Joers V, Weinshenker D, Tansey MG (2020) Transgenic Mice Expressing Human α-Synuclein in Noradrenergic Neurons Develop Locus Coeruleus Pathology and Nonmotor Features of Parkinson’s Disease. J Neurosci 40:7559–7576.

Carola V, D’Olimpio F, Brunamonti E, Mangia F, Renzi P (2002) Evaluation of the elevated plus-maze and open-field tests for the assessment of anxiety-related behaviour in inbred mice. Behavioural Brain Research 134:49–57.

Crum RM, Mojtabai R, Lazareck S, Bolton JM, Robinson J, Sareen J, Green KM, Stuart EA, La Flair L, Alvanzo AAH, Storr CL (2013) A Prospective Assessment of Reports of Drinking to Self-medicate Mood Symptoms with the Incidence and Persistence of Alcohol Dependence. JAMA Psychiatry 70:718–726.

Ferrazzoli D, Sica F, Sancesario G (2013) Sundowning Syndrome: A Possible Marker of Frailty in Alzheimer’s Disease? CNS & Neurological Disorders - Drug Targets 12:525–528.

Fish EW, Wieczorek LA, Rumple A, Suttie M, Moy SS, Hammond P, Parnell SE (2018) The Enduring Impact of Neurulation Stage Alcohol Exposure: A Combined Behavioral and Structural Neuroimaging Study in Adult Male and Female C57BL/6J Mice. Behav Brain Res 338:173–184.

Flurkey K, Currer JM, Harrison DE (2007), Mouse modes in aging research. Faculty Research 2000-2009. James G. Fox (ed.), American College of Laboratory Animal Medicine series (Elsevier, AP: Amsterdam; Boston).

Gallagher C, Radmall Z, O’Gara C, Burke T (2018) Anxiety and depression among patients with alcohol dependence: co-morbid or substance-related problems? Ir J Psychol Med 35:121–126.

Glöckner-Rist A, Lémenager T, Mann K, PREDICT Study Research Group (2013) Reward and relief craving tendencies in patients with alcohol use disorders: results from the PREDICT study. Addict Behav 38:1532–1540.

Gottesman RT, Stern Y (2019) Behavioral and Psychiatric Symptoms of Dementia and Rate of Decline in Alzheimer’s Disease. Front Pharmacol 10:1062.

Grucza RA, Sher KJ, Kerr WC, Krauss MJ, Lui CK, McDowell YE, Hartz S, Virdi G, Bierut LJ (2018) Trends in Adult Alcohol Use and Binge Drinking in the Early 21st-Century United States: A Meta-Analysis of 6 National Survey Series. Alcohol Clin Exp Res 42:1939–1950.

Gutwinski S, Schreiter S, Priller J, Henssler J, Wiers CE, Heinz A (2018) Drink and Think: Impact of Alcohol on Cognitive Functions and Dementia - Evidence of Dose-Related Effects. Pharmacopsychiatry 51:136–143.

Hart PC, Bergner CL, Smolinsky AN, Dufour BD, Egan RJ, LaPorte JL, Kalueff AV (2010) Experimental Models of Anxiety for Drug Discovery and Brain Research. In: Mouse Models for Drug Discovery: Methods and Protocols (Proetzel G, Wiles MV, eds), pp 299–321 Methods in Molecular Biology. Totowa, NJ: Humana Press. Available at: https://doi.org/10.1007/978-1-60761-058-8_18 [Accessed June 1, 2022].

Heinz A, Löber S, Georgi A, Wrase J, Hermann D, Rey E-R, Wellek S, Mann K (2003) Reward craving and withdrawal relief craving: assessment of different motivational pathways to alcohol intake. Alcohol Alcohol 38:35–39.

Heymann D, Stern Y, Cosentino S, Tatarina-Nulman O, Dorrejo JN, Gu Y (2016) The Association Between Alcohol Use and the Progression of Alzheimer’s Disease. Curr Alzheimer Res 13:1356–1362.

Hoffman JL, Faccidomo S, Kim M, Taylor SM, Agoglia AE, May AM, Smith EN, Wong LC, Hodge CW (2019) Alcohol drinking exacerbates neural and behavioral pathology in the 3xTg-AD mouse model of Alzheimer’s disease. In: International Review of Neurobiology, pp 169–230. Academic Press Inc.

Hwa LS, Chu A, Levinson SA, Kayyali TM, Debold JF, Miczek KA (2011) Persistent escalation of alcohol drinking in C57BL/6J mice with intermittent access to 20% ethanol. Alcoholism: Clinical and Experimental Research 35:1938–1947.

Ilomaki J, Jokanovic N, Tan E, Lonnroos E (2015) Alcohol Consumption, Dementia and Cognitive Decline: An Overview of Systematic Reviews. Current Clinical Pharmacology 10:204–212.

Kamal H, Tan GC, Ibrahim SF, Shaikh MF, Mohamed IN, Mohamed RMP, Hamid AA, Ugusman A, Kumar J (2020) Alcohol Use Disorder, Neurodegeneration, Alzheimer’s and Parkinson’s Disease: Interplay Between Oxidative Stress, Neuroimmune Response and Excitotoxicity. Front Cell Neurosci 14:282.

Kang SS, Liu X, Ahn EH, Xiang J, Manfredsson FP, Yang X, Luo HR, Liles LC, Weinshenker D, Ye K (2020) Norepinephrine metabolite DOPEGAL activates AEP and pathological Tau aggregation in locus coeruleus. J Clin Invest 130:422–437.

Keszycki RM, Fisher DW, Dong H (2019) The Hyperactivity–Impulsivity–Irritiability–Disinhibition–Aggression–Agitation Domain in Alzheimer’s Disease: Current Management and Future Directions. Front Pharmacol 10:1109.

Koob GF (2015) The dark side of emotion: The addiction perspective. European Journal of Pharmacology 753:73–87.

Koob GF, Le Moal M (1997) Drug abuse: hedonic homeostatic dysregulation. Science 278:52–58.

Koob GF, Le Moal M (2001) Drug addiction, dysregulation of reward, and allostasis. Neuropsychopharmacology 24:97–129.

Kosel F, Pelley JMS, Franklin TB (2020) Behavioural and psychological symptoms of dementia in mouse models of Alzheimer’s disease-related pathology. Neurosci Biobehav Rev 112:634–647.

Lacroix L, Spinelli S, Heidbreder CA, Feldon J (2000) Differential role of the medial and lateral prefrontal cortices in fear and anxiety. Behav Neurosci 114:1119–1130.

Lasagna-Reeves CA, de Haro M, Hao S, Park J, Rousseaux MWC, Al-Ramahi I, Jafar-Nejad P, Vilanova-Velez L, See L, De Maio A, Nitschke L, Wu Z, Troncoso JC, Westbrook TF, Tang J, Botas J, Zoghbi HY (2016) Reduction of Nuak1 Decreases Tau and Reverses Phenotypes in a Tauopathy Mouse Model. Neuron 92:407–418.

Leger M, Quiedeville A, Bouet V, Haelewyn B, Boulouard M, Schumann-Bard P, Freret T (2013) Object recognition test in mice. Nat Protoc 8:2531–2537.

Lister RG (1987) The use of a plus-maze to measure anxiety in the mouse. Psychopharmacology 92:180–185.

Lossos A, Reches A, Gal A, Newman JP, Soffer D, Gomori JM, Boher M, Ekstein D, Biran I, Meiner Z, Abramsky O, Rosenmann H (2003) Frontotemporal dementia and parkinsonism with the P301S tau gene mutation in a Jewish family. J Neurol 250:733–740.

Lowery-Gionta EG, Crowley NA, Bukalo O, Silverstein S, Holmes A, Kash TL (2018) Chronic stress dysregulates amygdalar output to the prefrontal cortex. Neuropharmacology 139:68–75.

Mann K, Roos CR, Hoffmann S, Nakovics H, Leménager T, Heinz A, Witkiewitz K (2018) Precision Medicine in Alcohol Dependence: A Controlled Trial Testing Pharmacotherapy Response Among Reward and Relief Drinking Phenotypes. Neuropsychopharmacology 43:891–899.

Masneuf S, Lowery-Gionta E, Colacicco G, Pleil KE, Li C, Crowley N, Flynn S, Holmes A, Kash T (2014) Glutamatergic mechanisms associated with stress-induced amygdala excitability and anxiety-related behavior. Neuropharmacology 85:190–197.

Menary KR, Kushner MG, Maurer E, Thuras P (2011) The prevalence and clinical implications of self-medication among individuals with anxiety disorders. J Anxiety Disord 25:335–339.

Morris HR, Khan MN, Janssen JC, Brown JM, Perez-Tur J, Baker M, Ozansoy M, Hardy J, Hutton M, Wood NW, Lees AJ, Revesz T, Lantos P, Rossor MN (2001) The genetic and pathological classification of familial frontotemporal dementia. Arch Neurol 58:1813–1816.

Mozhui K, Karlsson R-M, Kash TL, Ihne J, Norcross M, Patel S, Farrell MR, Hill EE, Graybeal C, Martin KP, Camp M, Fitzgerald PJ, Ciobanu DC, Sprengel R, Mishina M, Wellman CL, Winder DG, Williams RW, Holmes A (2010) Strain Differences in Stress Responsivity Are Associated with Divergent Amygdala Gene Expression and Glutamate-Mediated Neuronal Excitability. J Neurosci 30:5357–5367.

Osborn JA, Kim CK, Steiger J, Weinberg J (1998) Prenatal ethanol exposure differentially alters behavior in males and females on the elevated plus maze. Alcohol Clin Exp Res 22:685–696.

Panza F, Frisardi V, Seripa D, Logroscino G, Santamato A, Imbimbo BP, Scafato E, Pilotto A, Solfrizzi V (2012) Alcohol consumption in mild cognitive impairment and dementia: harmful or neuroprotective? Int J Geriatr Psychiatry 27:1218–1238.

Pentkowski NS, Rogge-Obando KK, Donaldson TN, Bouquin SJ, Clark BJ (2021) Anxiety and Alzheimer’s disease: Behavioral analysis and neural basis in rodent models of Alzheimer’s-related neuropathology. Neuroscience & Biobehavioral Reviews 127:647–658.

Prut L, Belzung C (2003) The open field as a paradigm to measure the effects of drugs on anxiety-like behaviors: a review. Eur J Pharmacol 463:3–33.

Rehm J, Hasan OSM, Black SE, Shield KD, Schwarzinger M (2019) Alcohol use and dementia: a systematic scoping review. Alzheimers Res Ther 11:1.

Ridley NJ, Draper B, Withall A (2013) Alcohol-related dementia: an update of the evidence. Alzheimers Res Ther 5:3.

Rotermund C, Reolon GK, Leixner S, Boden C, Bilbao A, Kahle PJ (2017) Enhanced motivation to alcohol in transgenic mice expressing human α-synuclein. J Neurochem 143:294–305.

Scattoni ML, Gasparini L, Alleva E, Goedert M, Calamandrei G, Spillantini MG (2010) Early behavioural markers of disease in P301S tau transgenic mice. Behavioural Brain Research 208:250–257.

Sperfeld AD, Collatz MB, Baier H, Palmbach M, Storch A, Schwarz J, Tatsch K, Reske S, Joosse M, Heutink P, Ludolph AC (1999) FTDP-17: an early-onset phenotype with parkinsonism and epileptic seizures caused by a novel mutation. Ann Neurol 46:708–715.

Takeuchi H, Iba M, Inoue H, Higuchi M, Takao K, Tsukita K, Karatsu Y, Iwamoto Y, Miyakawa T, Suhara T, Trojanowski JQ, Lee VM-Y, Takahashi R (2011) P301S Mutant Human Tau Transgenic Mice Manifest Early Symptoms of Human Tauopathies with Dementia and Altered Sensorimotor Gating Bush AI, ed. PLoS ONE 6:e21050.

Todd WD (2020) Potential Pathways for Circadian Dysfunction and Sundowning-Related Behavioral Aggression in Alzheimer’s Disease and Related Dementias. Frontiers in Neuroscience 14 Available at: https://pubmed.ncbi.nlm.nih.gov/33013301/ [Accessed April 27, 2021].

Turner S, Mota N, Bolton J, Sareen J (2018) Self-medication with alcohol or drugs for mood and anxiety disorders: A narrative review of the epidemiological literature. Depression and Anxiety 35:851–860.

Tyas SL (2001) Alcohol use and the risk of developing Alzheimer’s disease. Alcohol Res Health 25:299–306.

Watt G, Przybyla M, Zak V, van Eersel J, Ittner A, Ittner LM, Karl T (2020) Novel Behavioural Characteristics of Male Human P301S Mutant Tau Transgenic Mice – A Model for Tauopathy. Neuroscience 431:166–175.

Werber E, Klein C, Grünfeld J, Rabey JM (2003) Phenotypic presentation of frontotemporal dementia with Parkinsonism-chromosome 17 type P301S in a patient of Jewish-Algerian origin. Mov Disord 18:595–598.

White AM (2020) Gender Differences in the Epidemiology of Alcohol Use and Related Harms in the United States. Alcohol Res 40:01.

Wilson SR, Knowles SB, Huang Q, Fink A (2014) The Prevalence of Harmful and Hazardous Alcohol Consumption in Older U.S. Adults: Data from the 2005–2008 National Health and Nutrition Examination Survey (NHANES). J Gen Intern Med 29:312–319.

Wise RA, Koob GF (2014) The development and maintenance of drug addiction. Neuropsychopharmacology 39:254–262.

Xu H, Rösler TW, Carlsson T, Andrade A de, Bruch J, Höllerhage M, Oertel WH, Höglinger GU (2014) Memory deficits correlate with tau and spine pathology in P301S MAPT transgenic mice. Neuropathology and Applied Neurobiology 40:833–843.

Xu W, Wang H, Wan Y, Tan C, Li J, Tan L, Yu J-T (2017) Alcohol consumption and dementia risk: a dose-response meta-analysis of prospective studies. Eur J Epidemiol 32:31–42.

Yoshiyama Y, Higuchi M, Zhang B, Huang SM, Iwata N, Saido TCC, Maeda J, Suhara T, Trojanowski JQ, Lee VMY (2007) Synapse Loss and Microglial Activation Precede Tangles in a P301S Tauopathy Mouse Model. Neuron 53:337–351.

Flurkey, K; Currer, J M.; and Harrison, D E., “Mouse models in aging research.” (2007). Faculty Research 2000 - 2009. 1685. https://mouseion.jax.org/stfb2000_2009/1685

